# Cell type-specific translational regulation by human DUS enzymes

**DOI:** 10.1101/2023.11.03.565399

**Authors:** Nathan J. Yu, Wei Dai, Ang Li, Muhan He, Ralph E. Kleiner

**Affiliations:** Department of Chemistry, Princeton University, Princeton, NJ, USA 08544

## Abstract

Dihydrouridine is an abundant and conserved modified nucleoside present on tRNA, but characterization and functional studies of modification sites and associated DUS writer enzymes in mammals is lacking. Here we use a chemical probing strategy, RNABPP-PS, to identify 5-chlorouridine as an activity-based probe for human DUS enzymes. We map D modifications using RNA-protein crosslinking and chemical transformation and mutational profiling to reveal D modification sites on human tRNAs. Further, we knock out individual DUS genes in two human cell lines to investigate regulation of tRNA expression levels and codon-specific translation. We show that whereas D modifications are present across most tRNA species, loss of D only perturbs the translational function of a subset of tRNAs in a cell type-specific manner. Our work provides powerful chemical strategies for investigating D and DUS enzymes in diverse biological systems and provides insight into the role of a ubiquitous tRNA modification in translational regulation.

## INTRODUCTION

Post-transcriptional modifications on RNA play an important role in biological processes and their dysregulation is associated with disease^1, 2^. To date, over 150 different chemical modifications^3^ have been identified on cellular RNA, but functional characterization and transcriptome-wide mapping data is lacking for many modifications. Among RNA species, tRNAs are the most heavily modified^4, 5^,and tRNA modifications are highly conserved and involved in regulating tRNA stability and folding, charging by aminoacyl-tRNA synthetases enzymes, and codon-anticodon pairing. Defects in tRNA modifying enzymes have been implicated in a host of human diseases known as tRNA modopathies^6^ and dysregulation of tRNA modifications can drive the translation of oncogenic proteins and cellular transformation^7^. Considerable effort has been focused on the study of tRNA modifications, yet the biological function of many abundant and evolutionarily conserved tRNA modifications still remains enigmatic. Dihydrouridine (D) is among the most abundant tRNA modifications in biology^8^. Whereas D was originally identified in the eponymous tRNA “D arm” by Holley^9^, the enzymes responsible for its installation, dihydrouridine synthases (DUS), were not characterized until decades later in *E. coli*^10^ and *S. cerevisiae*^11^. Eukaryotes possess four DUS enzymes and the specificity of the yeast proteins for distinct tRNA sites was established by Phizicky and co-workers^12^ using primer extension and microarray analysis. However, deletion of DUS enzymes in yeast has little effect on growth^12^. Interestingly, two recent studies identified the presence of D in *S. pombe*^13^ and *S. cerevisiae*^14^ mRNA, and Finet *et al.*^13^ demonstrated that dihydrouridylation on α-tubulin mRNA in *S. pombe* affects protein expression and meiotic chromosome segregation. These works expand the scope of D modification across the eukaryotic transcriptome, but the functions of tRNA D modifications, which are the major substrates of DUS enzymes, are not well understood. Further, elevated levels of DUS proteins and D have been associated with malignancy and poor patient outcomes^15-18^, but biochemical and cell biological studies of human DUS enzymes are lacking.

A major obstacle to the study of D is the challenge in characterizing modification levels across diverse RNA species and assigning relevant writer enzymes. Reported chemical mapping strategies rely upon reverse transcription termination events^12-14^, which are not well suited to mapping tRNA D sites due to the close proximity of D modifications and high density of other native modifications. Further, they do not enable enrichment of low abundance sites. Previously we developed an activity-based strategy for discovering RNA modifying enzymes and characterizing their RNA substrates. Our approach, RNABPP^19, 20^, uses metabolic labeling with modified nucleosides to induce mechanism-based crosslinking between RNA modifying enzymes and their substrate RNAs for proteomic or transcriptomic analysis. Our prior work^19^ demonstrated that 5-fluorouridine (5-FUrd) is a mechanism-based probe for human DUS3L, and we exploited our covalent warhead to map DUS3L substrates to the variable loop of human tRNA. We also showed that DUS3L KO cells are compromised in cell proliferation and protein translation efficiency. Interestingly, we did not identify mechanism-based crosslinks between 5-FUrd-labeled RNA and other human DUS enzyme homologs, despite likely conservation in catalytic mechanism^21^. We also did not investigate further the basis of the observed translational defects.

Here, we develop RNABPP-PS, an activity-based RNA modifying enzyme probing strategy relying upon metabolic labeling with modified nucleosides and phase separation-based protein-RNA enrichment. In contrast to our RNABPP strategy that relied upon oligo(dT) enrichment, RNABPP-PS is not biased towards the capture of mRNA modifying enzymes. We apply RNABPP-PS with diverse 5-halopyrimidine nucleosides to demonstrate that 5-chlorouridine (5-ClUrd) is a general mechanism-based probe for human DUS enzymes, and profile DUS1L and DUS2L substrates using catalytic crosslinking and RNA sequencing. We further map D sites across human tRNAs in HEK293T and HeLa cells using chemical reactivity coupled with high-throughput sequencing and mutational profiling. Finally, we explore the effect of DUS KO on cell proliferation, tRNA stability, and codon-specific translation, to show that human DUS proteins regulate translation and tRNA expression levels in a cell type-specific manner. Taken together, our study provides a suite of strategies for studying DUS enzymes and mapping D sites across the transcriptome and reveal new insights into the role of these proteins and their associated modifications in tRNA biology and translational regulation.

## RESULTS

### Phase-separation based enrichment of cellular RNA modifying enzymes

To globally capture RNA modifying enzymes for proteomic analysis, we needed a method to isolate and enrich these proteins from biological samples. We previously developed RNABPP^19^, which combines metabolically incorporated reactive nucleosides with oligo(dT)-based enrichment of crosslinked RNA-protein complexes. The reliance upon oligo(dT)-enrichment biases RNABPP towards enzymes that modify polyadenylated mRNA, therefore alternative methods are required to broadly enrich RNA modifying enzymes irrespective of the identity of their RNA substrate. One such approach is organic-aqueous phase separation^22-24^, which has been used to isolate UV-crosslinked RNA-protein complexes based on their propensity to partition into an insoluble “interphase” (Fig. 1a). Whereas this method has been primarily applied to characterize RNA-binding proteins that interact with mRNA, we hypothesized that RNA-modifying enzymes crosslinked to their RNA substrates (which can range in size over several orders of magnitude) using modified nucleoside probes would also be amenable to phase separation-based enrichment and proteomic analysis.

**Figure 1.**
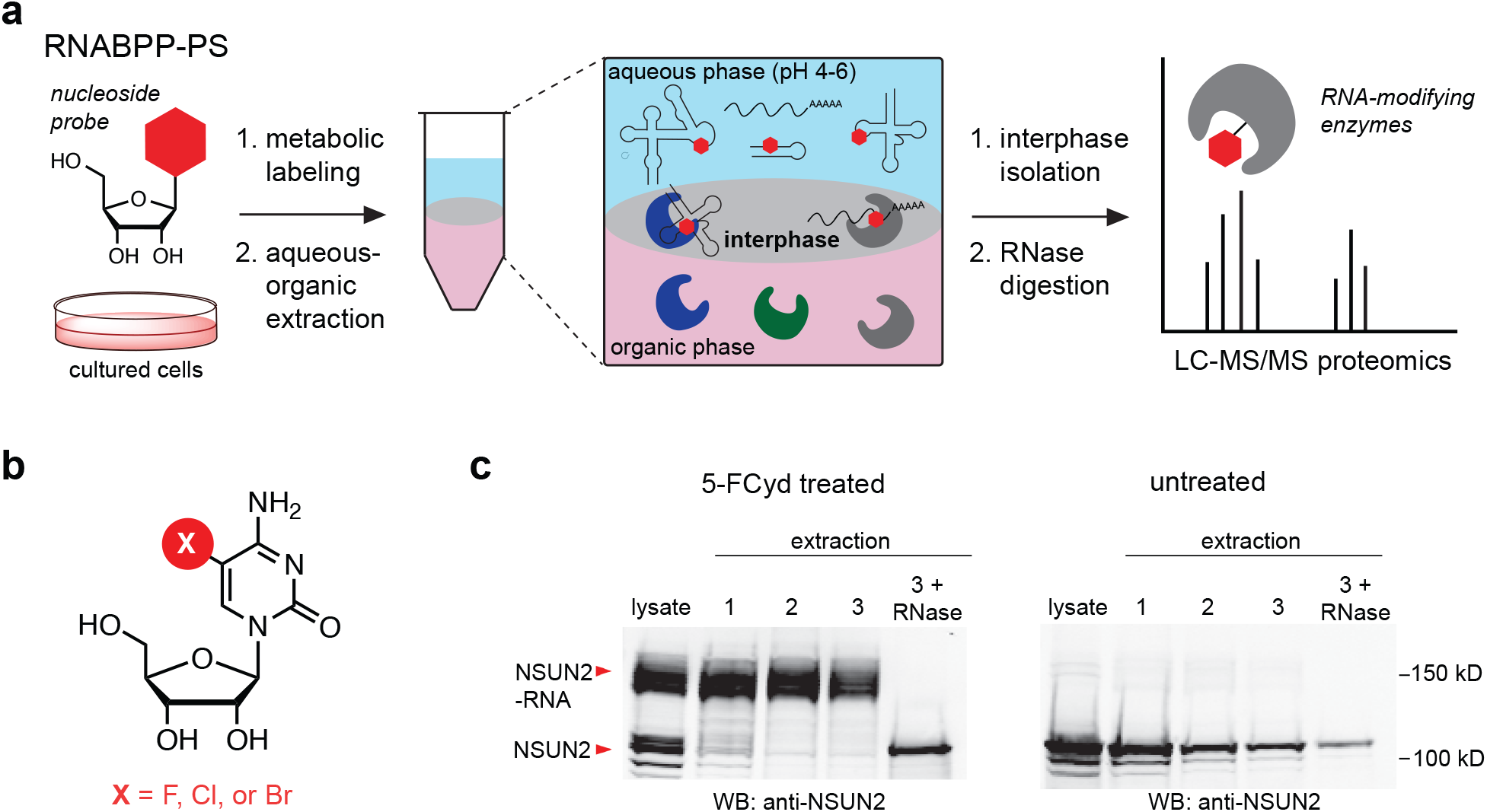
Phase separation-based enrichment of catalytically crosslinked RNA-modifying enzymes. **(a)** Schematic of RNABPP-PS method. RNA-protein complexes are crosslinked through metabolic incorporation of a mechanism-based nucleoside probe and enriched by aqueous-organic phase separation. Reactive proteins are then identified by mass spectrometry-based proteomics. **(b)** Chemical structure of 5-halopyrimidine activity-based nucleosides used in this work. **(c)** Western blot analysis of NSUN2-RNA covalent complexes present in interphase samples after metabolic labeling of RNA with 5-FCyd. Cells were treated with 10 µM 5-FCyd for 16 h or untreated and then subjected to repeated cycles of aqueous-organic extraction, followed by RNAse digestion.

To validate this approach, we employed metabolic labeling with 5-fluorocytidine (5-FCyd) (Fig. 1b), which we showed previously^19^ is an efficient activity-based probe for RNA C5-methyltransferases as well as dihydrouridine synthases (DUS), through deamination to 5-FUrd. We tested whether organic-aqueous extraction would enrich RNA-protein crosslinks formed between candidate proteins NSUN2 and DUS3L, which we had isolated in previous RNABPP studies with 5-FCyd and oligo(dT)-pulldown. In brief, cells were treated with 5-FCyd and subjected to lysis with a mixture of acidic guanidinium isothiocyanate/phenol/chloroform (i.e. TRIzol or TRI reagent). The insoluble interphase was isolated, subjected to repeated rounds of organic-aqueous extraction, and proteins were released with RNase A/T1 digestion. In 5-FCyd treated cells, western blot analysis of NSUN2 and DUS3L in cellular lysate and extracted interphase samples demonstrated clear formation of a higher running RNA-protein species that selectively partitioned into the interphase (Fig. 1c, Supplementary Fig. 1). In contrast, free protein was present in lysate but was not retained in the interphase after sequential organic-aqueous extractions. Using this approach, we recovered 3-4-fold more NSUN2 or DUS3L protein in 5-FCyd-treated cells as compared to non-treated control samples, indicating that mechanism-based nucleoside probes and phase separation (i.e. RNABPP-PS) can be applied to globally profile RNA modifying enzymes in living cells.

### Proteomic profiling of human RNA modifying enzymes with C5-halopyrimidines

Having validated the RNABPP-PS approach, we applied our reactivity-based method to profile RNA modifying enzymes in human cells. Due to the relative simplicity of the phase separation-based enrichment protocol and low sample input requirement, we decided to interrogate multiple C5-modified halopyrimidine nucleosides (Fig. 1b) to broadly capture RNA modifying enzymes. First, we established that metabolic labeling of cellular RNA proceeds efficiently with 5-chlorocytidine (5-ClCyd) and 5-bromocytidine (5-BrCyd). Whereas these nucleosides are larger than 5-FCyd and less efficient substrates for enzymes in the pyrimidine salvage pathway^25^, increasing the metabolic labeling concentration (250-500 µM) achieved similar levels of RNA incorporation (∼0.5-1.0% of C) as with 10 µM 5-FCyd (Supplementary Fig. 1a, Supplementary Table 1), without overt cytotoxicity. Additionally, all three cytidine nucleosides were deaminated at comparable rates to the corresponding modified uridine derivatives.

Next, we prepared RNABPP-PS samples from cells grown in the presence of 5-FCyd, 5-ClCyd, or normal medium (control) for quantitative proteomic analysis. Independent biological replicates (three for all conditions) were labeled with tandem mass tags (TMT)^26^, combined, and measured in one LC-MS/MS run, similar to our RNABPP pipeline^19^. In the 5-FCyd RNABPP-PS experiment (Fig. 2a, Supplementary Datafile 1), the top four hits with highest fold change were all known RNA C5-methyltransferase enzymes. These include three members of the NSUN family – NSUN2, NSUN5, and NSUN6 – which generate m^5^C on various RNA substrates including mRNA, rRNA, and tRNA^27^, and TRMT2A, an RNA 5-methyluridine (m^5^U) methyltransferase that primarily installs m^5^U54 on tRNA^28^. We also found DUS3L, the dihydrouridine synthase that we used to validate our method. As expected, 5-FCyd RNABPP-PS recovered many of the same proteins that were identified by 5-FCyd RNABPP with oligo(dT)-based enrichment, thereby validating the approach for profiling RNA m^5^U and m^5^C methyltransferases and dihydrouridine synthases. One notable exception was NSUN6, which was readily captured through RNABPP-PS but not identified in our earlier study. Additional proteins that were enriched (>2-fold) upon 5-FCyd treatment and interphase extraction – SDC1/3, COQ9, GAR1, and DDX46 – do not have obvious RNA modification domains and were likely enriched due to physicochemical properties that favor partitioning into the interphase and/or upregulation upon 5-FCyd treatment.

**Figure 2.**
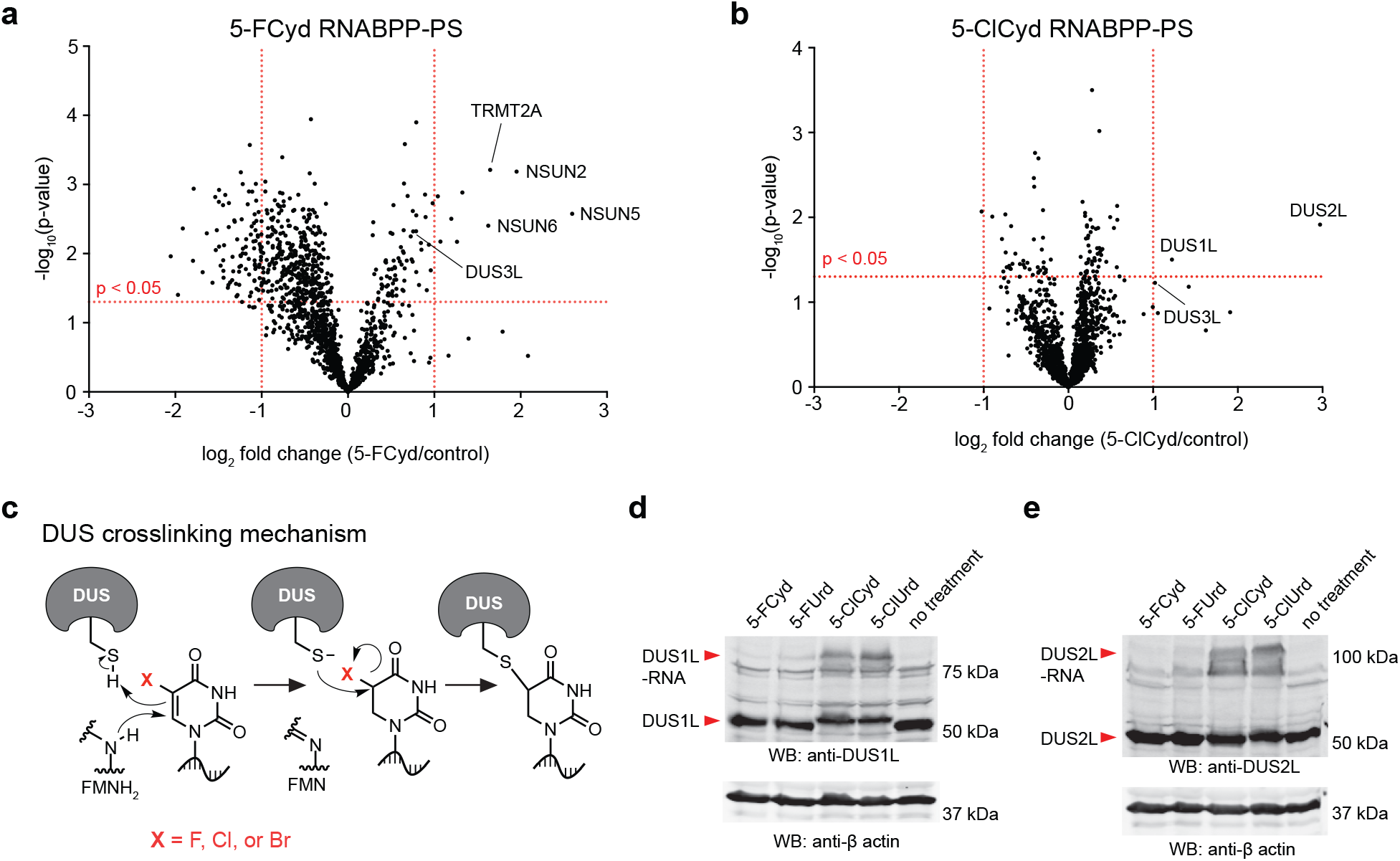
Proteomic profiling of 5-FCyd- and 5-ClCyd-reactive proteins with RNABPP-PS. **(a)** Volcano plot of 5-FCyd-reactive proteins. Known RNA modifying enzymes are annotated. **(b)** Volcano plot of 5-ClCyd-reactive proteins. Known RNA modifying enzymes are annotated. **(c)** Proposed mechanism of DUS crosslinking by C5-halogenated uracil bases. **(d)-(e)** Western blot analysis of DUS1L and DUS2L protein-RNA crosslinking after metabolic labeling with a panel of C5-halogenated pyrimidines.

Analysis of 5-ClCyd-enriched proteins revealed a much narrower scope of enzyme reactivity than 5-FCyd (Fig. 2b). We identified only two proteins showing significant fold change >2.0: DUS1L and DUS2L. DUS3L was also enriched although just below the p<0.05 threshold. All three proteins are mammalian homologs of yeast tRNA dihydrouridine synthases (DUS)^12^, suggesting that 5-ClCyd (likely through deamination to 5-ClUrd) is a selective and general probe for these enzymes. This stands in contrast to 5-FUrd, which reacts primarily with DUS3L. Based on our proposed RNA-protein crosslinking mechanism (Fig. 2c)^19^, it is likely that substitution of F for Cl accelerates nucleophilic displacement by the conserved catalytic Cys residue in DUS enzymes^21,29^. To validate mechanism-based crosslinking with 5-chloropyrimidines, we performed western blot analysis of DUS1L and DUS2L in cellular lysate after nucleoside treatment. As expected, treatment of cells with 5-FCyd or 5-FUrd did not generate an observable crosslink, whereas metabolic labeling with 5-ClCyd or 5-ClUrd produced a slower migrating band (Fig. 2d, 2e, Supplementary Fig. 2), consistent with the formation of a covalent RNA-protein complex. These findings support our RNABPP-PS proteomic data and establish that 5-chloropyrimidines are efficient activity-based nucleoside probes for mammalian DUS enzymes.

### Human DUS proteins regulate cell proliferation and protein translation

To investigate the role of DUS enzymes in mammalian biology, we generated DUS KO cell lines using CRISPR/Cas technology^30^ in HEK 293T as the genetic background. Previously, we isolated and characterized DUS3L KO cells^19^ and identified defects in cell proliferation and protein translation. In this study, we generated HEK293T clones containing individual knockout of DUS1L or DUS2L, double knockout of DUS1L and DUS2L, and quadruple knockout of DUS1L-DUS4L (Supplementary Fig. 3-8, Supplementary Table 2). Cell lines were confirmed by western blot and genomic sequencing, and we measured D levels in total RNA and fractionated small RNA (17 to 300 nt) using nucleoside LC-QQQ-MS (Fig. 3a, 3b, Supplementary Tables 3, 4). D levels in total RNA accounted for ∼1.0% of U residues and ∼10% of U in small RNA, consistent with D residing primarily on tRNA. In DUS KO cells, we observed partial reductions in D levels in individual KO cells, additive reductions in the double DUS1L/2L KO, and complete loss of D in the quadruple DUS KO cell line. Our data indicates that the four mammalian DUS enzymes are responsible for all D modifications on RNA and suggest non-redundant substrate sites.

**Figure 3.**
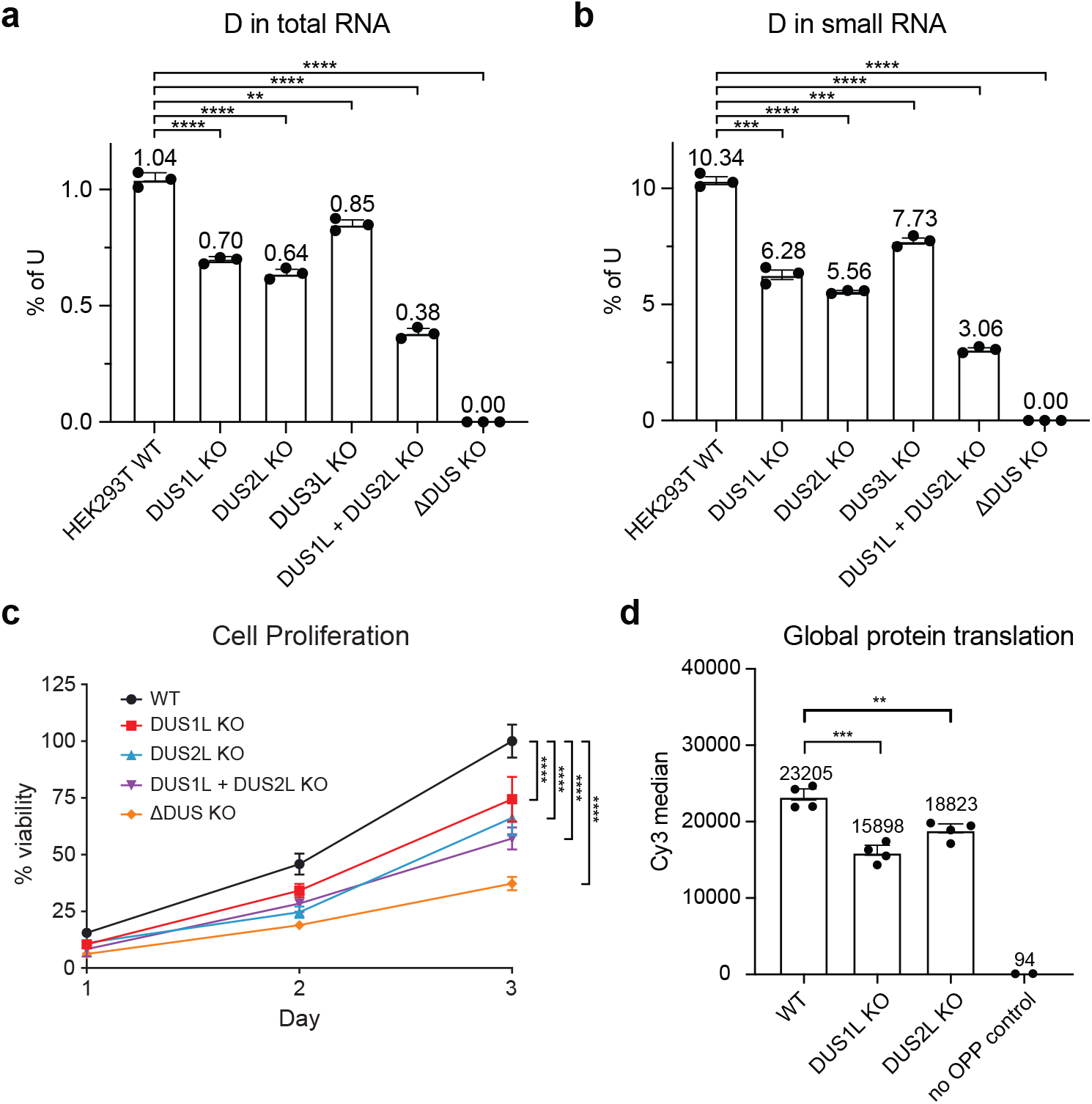
Characterization of HEK293T DUS KO cell lines. **(a)-(b)** Nucleoside LC-QQQ-MS analysis of D levels in total RNA and small RNA. Three independent biological replicates were analyzed per cell line. Data bars are representative of mean values ± s.e.m. Unpaired t-test (two-tailed) were used to evaluate statistical significance. *P*-values: WT vs. DUS1L KO in total RNA, 0.000067; WT vs. DUS2L KO in total RNA, 0.000062; WT vs. DUS3L KO in total RNA, 0.0012; WT vs. DUS1L + DUS2L KO in total RNA, 0.0000088; WT vs. ΔDUS in total RNA, 0.00000058; WT vs. DUS1L KO in small RNA, 0.00011; WT vs. DUS2L KO in small RNA, 0.000011; WT vs. DUS3L KO in small RNA, 0.00028; WT vs. DUS1L + DUS2L KO in small RNA, 0.0000026; WT vs. ΔDUS in total RNA, 0.00000044. **(c)** Cell proliferation analysis of HEK293T DUS KO cell lines. Four independent biological replicates were analyzed. Data represent mean values ± s.e.m. Unpaired t-tests (two-tailed) were performed to evaluate the statistical significance. *P*-values: WT vs. DUS1L KO, 0.000042; WT vs. DUS2L KO, 0.00000035; WT vs. DUS1L + DUS2L KO, 0.0000000018; WT vs. ΔDUS, 0.00000000000063. **(d)** Global protein translation efficiency in HEK293T WT, DUS1L KO, and DUS2L KO cells. OP-puro incorporation in nascent proteins was measured by flow cytometry. The median fluorescence intensity and s.e.m. values from four independent biological replicates (50,000 cells analyzed per sample, three technical replicates per sample) are shown. Unpaired t-test (two-tailed) was performed to measure statistical significance. *P-values*: WT vs. DUS1L, 0.00029; WT vs. DUS2L, 0.0036.

We further studied cell viability and protein translation in our HEK293T DUS KO cell lines. Compared to WT, we observed 30% and 40% reduction in viability after 3 d in DUS1L and DUS2L KO lines, respectively (Fig. 3c, Supplementary Table 5), and 50% reduction in the DUS1L/2L double KO. Finally, the quadruple DUS KO cell line exhibited the poorest viability and failed to proliferate when passaged at low confluence. To study protein translation, we measured *O-* propargyl-puromycin (OP-puro)^31^ incorporation into nascent proteins. We observed 31% and 19% reduction in OP-puro labeling (Fig. 3d, Supplementary Table 6, Supplementary Fig. 9) in DUS1L KO and DUS2L KO strains, respectively. These results indicate general impairment in protein translation efficiency and cell proliferation resulting from DUS1L or DUS2L knockdown, similar to our previous findings with DUS3L^19^, and additive effects when depleting multiple DUS proteins in the same cell line.

### Mapping DUS1L- and DUS2L-mediated D sites using 5-ClUrd-iCLIP

The substrates of DUS1L and DUS2L have not been characterized in human cells, therefore we adapted the iCLIP method^32^ to map DUS1L- and DUS2L-dependent D sites by combining 5-ClUrd metabolic labeling and crosslinking with high-throughput RNA sequencing. Previously we used this approach with 5-FUrd to establish the cellular substrates of DUS3L^19^. Briefly, cells expressing epitope-tagged DUS1L or DUS2L (Supplementary Fig. 10) were fed 5-ClUrd and DUS proteins were immunoprecipitated together with covalently linked RNA (Fig. 4a). As expected, DUS proteins immunoprecipitated from 5-ClUrd-treated cells contained slower migrating species that contained RNA, indicative of efficient mechanism-based crosslinking (Fig. 4b). We generated iCLIP libraries from 5-ClUrd-crosslinked RNA for DUS1L and DUS2L following literature precedent (Supplementary Fig. 11)^19^. Control libraries were prepared in parallel from untreated cells, and we performed bioinformatic analysis to quantify reverse transcription termination events (i.e. “RT stops”) enriched in the 5-ClUrd samples (corresponding to D sites) relative to control.

**Figure 4.**
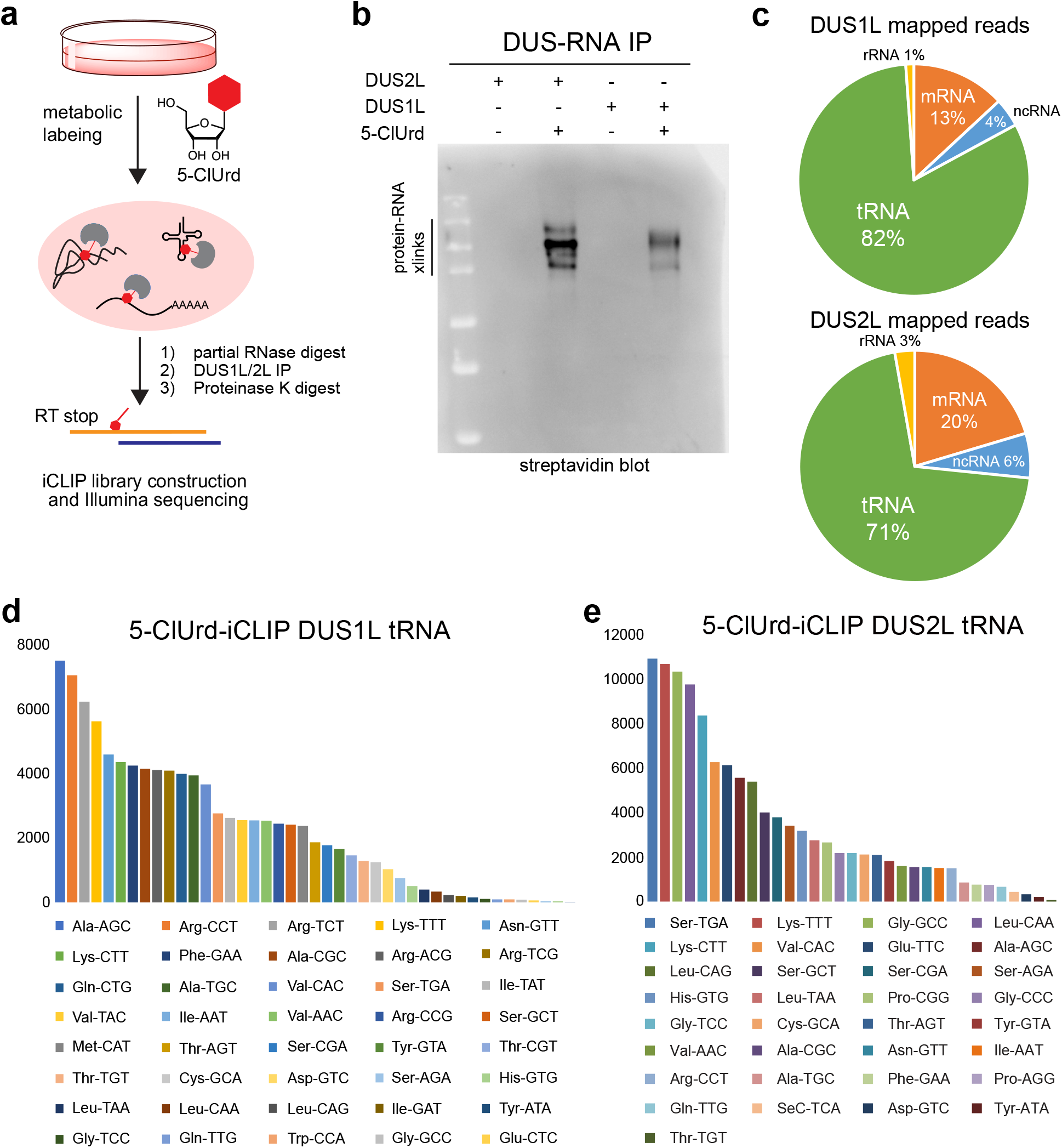
5-ClUrd-iCLIP analysis of DUS1L and DUS2L substrates. **(a)** Schematic of 5-ClUrd-iCLIP workflow; IP, immunoprecipitation; RT, reverse transcription. **(b)** Demonstration of RNA-protein crosslinking in DUS2L and DUS1L iCLIP samples. Crosslinked RNA was detected by streptavidin western blot following immunoprecipitation, RNA fragmentation, and ligation of a biotinylated adaptor. Experiment was repeated three times independently with similar results. **(c)** Composition of uniquely mapped reads in 5-ClUrd-iCLIP experiments for DUS1L and DUS2L. **(d)-(e)** Abundance of tRNA anticodon species found in 5-ClUrd-iCLIP for DUS1L and DUS2L. tRNAs are sorted by reads per million (RPM) of mapped reads.

70-80% of uniquely mapped reads in the 5-ClUrd-treated DUS1L and DUS2L samples mapped to tRNA, indicating that tRNAs are the primary substrates of these enzymes (Fig. 4c, Supplementary Datafile 2). This is consistent with our LC-QQQ-MS data as well as the substrate preferences of the yeast homologues Dus1p and Dus2p^11, 12^. We rank ordered tRNA isodecoders based on abundance and observed enrichment of distinct tRNAs in the DUS1L and DUS2L samples, indicating that these enzymes have different preferred tRNA substrates (Fig. 4d, 4e). For example, among the 43 tRNA isodecoder families detected for DUS1L, the most abundant were Ala-AGC, Arg-CCT, and Arg-TCT, whereas the top tRNAs among the 18 tRNA isodecoder families detected for DUS2L were Lys-TTT, Val-CAC, and Glu-TTC (Fig. 4d and 4e). We next analyzed the positional distribution of RT stop peaks from all tRNA aligned reads. We restricted our analysis to RT stops occurring at or adjacent to U residues to focus on RT stops induced by 5-ClUrd-mediated crosslinking events. Although multiple peaks were detected across the length of the tRNA (most likely due to endogenous tRNA modifications), we observed a primary peak at position 15 in the DUS1L sample (Supplementary Fig. 11). We also found major peaks at positions 19-20 in DUS2L, however the largest peak occurred at position 26, likely due to the native m^2^_2_G modification found at this position^33^. In light of the known substrate preferences of the corresponding yeast homologues (i.e. Dus1p modifies U16/17 and Dus2p modifies U20)^11, 12^, the uncertainty in RT stop position relative to the crosslinked adduct, and the confounding effect of native tRNA modifications on RT, we conclude that DUS1L and DUS2L likely modify U16/17 and U20 in the tRNA D loop, respectively.

### Mapping tRNA D sites with NaBH_4_-based mutational sequencing

Due to the challenge in assigning precise D modification sites on tRNA from the RT stop-based iCLIP analysis, we decided to pursue a mapping strategy relying upon RT misincorporation events rather than RT stops. Whereas D does not efficiently induce misincorporation^13^, we hypothesized that chemoselective reduction of D to tetrahydrouridine (THU) with NaBH_4_^34^ would generate a distinctive RT signature since the Watson-Crick face of THU is distinct from U or D (Fig. 5a). A similar strategy has been used for sequencing ac^4^C through borohydride-mediated reduction to tetrahydrocytidine^35^. NaBH_4_-based sequencing of D has been used in two recent methods, Rho-seq and D-seq^13, 14^, however both rely upon NaBH_4_-induced RT stops rather than misincorporation making them challenging to apply to tRNA due to the abundance of endogenous RT-stop inducing tRNA modifications. To pilot our method, we performed RT-PCR analysis of tRNA-Val-CAC from WT HEK293T cells and matched DUS3L KO cells. We identified D47 on Val-CAC as a DUS3L substrate in our earlier work^19^. As expected, targeted sequencing around the D47 site did not generate RT mutations in tRNA analyzed from WT or DUS3L KO cells, however we found ∼30% T-to-C mutation specifically at position 47 when RNA from WT cells was treated with NaBH_4_ before RT (Supplementary Fig. 12). In contrast, NaBH_4_-treated RNA from DUS3L KO cells lacked any mutation at U47 indicating that the NaBH_4_-induced mutational signature was specific to D. Encouraged by this result, we generated high-throughput sequencing libraries from total tRNA isolated from WT HEK293T or DUS3L KO cells by adapting the ARM-seq/DM-seq methods^36, 37^ and analyzed NaBH_4_-dependent mutational events across all identified tRNA species using VarScan variant detection software^38^. In tRNA from WT cells, we measured 432 NaBH_4_- dependent mutational events occurring at U residues across 197 tRNA species. These mutations largely occur near known D positions – 16,17, 20, 20a, 20b, and 47 (Fig. 5b, Supplementary Fig. 13, Supplementary Datafile 3); gratifyingly, tRNA from DUS3L KO cells showed a dramatic reduction in mutational events at U47 (Fig. 5b, Supplementary Fig. 14, Supplementary Datafile 3) whereas mutations at D loop positions were largely unchanged, demonstrating the specificity of our method.

**Figure 5.**
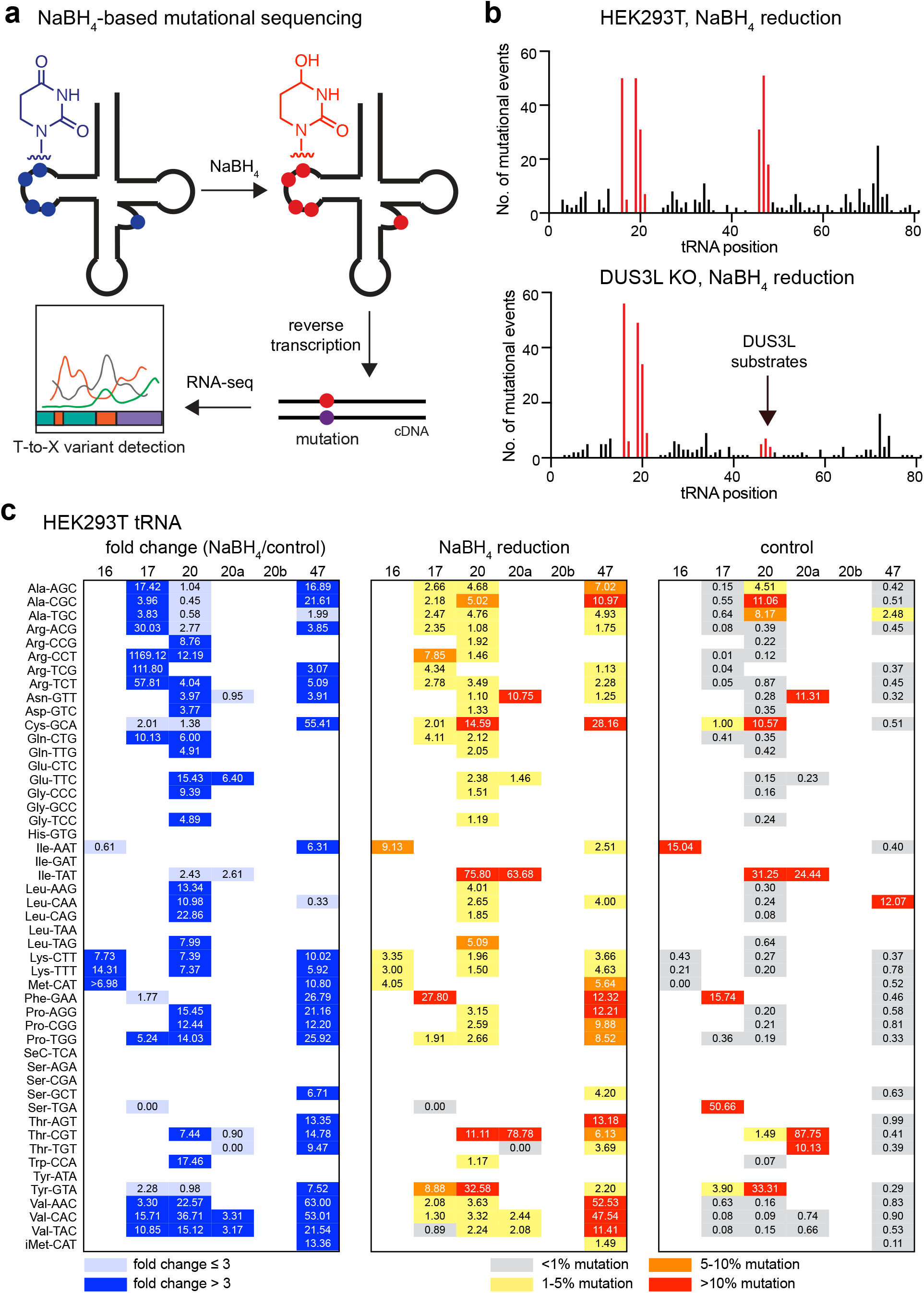
NaBH_4_-based mutational profiling of D sites in tRNA. **(a)** Scheme for NaBH_4_ reduction of D and misincorporation during RT. **(b)** NaBH_4_-dependent mutational events across all sequenced tRNAs from HEK293T and DUS3L KO cells. **(c)** Misincorporation rates at putative tRNA D sites in NaBH_4_-treated and untreated RNA from HEK293T cells.

Next, we analyzed individual mutational events at predicted D positions (using the Sprinzl tRNA position numbering system^39^) in NaBH_4_-treated WT RNA and the untreated control for all tRNA isodecoder families in our data. We collapsed isodecoder-specific data according to tRNA anticodon sequence and found 66 potential D modification sites that showed 3-fold or higher mutation rate in the NaBH_4_-treated sample as compared to the control, with the majority exhibiting 10-100-fold difference in mutation rate (Fig. 5c). Of these, 12 sites were found at position 17, 24 sites were found at position 20, and 24 sites were found at position 47. We identified only 3 modification sites at position 16 and 3 sites at position 20a; no sites were found at position 20b. Several sites at position 20/20a exhibited high mutation rates (>10%) in both NaBH_4_-treated and control samples (i.e. position 20a on tRNA-Asn-GTT, position 20 on tRNA-Cys-GCA, positions 20/20a on tRNA-Ile-TAT, position 20a on tRNA-Thr-CGT, and position 20 on tRNA-Tyr-GTA) – 4 of these sites are known to contain the acp^3^U modification that can cause RT misincorporation^40^. Inspection of the prevalence of U residues in tRNA sequences at potential D modification sites indicated that most tRNAs that contain U residues at these positions are represented in our data and are therefore substrates for dihydrouridylation. Notable exceptions are tRNA-Ser isoacceptors, among which we found only a single D modification, and tRNA-SeC and tRNA-His in which no NaBH_4_-dependent mutations were found.

Interestingly, whereas the relative change (i.e. fold change) in mutation induced by NaBH_4_ was similar across different tRNA D positions, the mutation rate in the NaBH_4_-treated sample showed considerable variation (Fig. 5c). In particular, positions in the D arm (i.e. 16, 17, 20, 20a, and 20b) consistently showed more modest misincorporation frequencies ranging between 1-5%, with the exception of the high mutation rates that likely result from acp^3^U modifications at 20/20a. In contrast, several D47 modifications in the variable loop result in ∼10-50% misincorporation frequency upon NaBH_4_ reduction, including D47 sites in tRNA-Val, tRNA-Thr, tRNA-Pro, tRNA-Phe, tRNA-Cys, and tRNA-Ala isoacceptors. While comprehensive data on the stoichiometry of D modification sites on human tRNAs is unavailable, we propose that the NaBH_4_-dependent mutational signature reflects not only the modification abundance but is also a function of sequence/structure-dependent differences in NaBH_4_ reactivity and misincorporation rates during reverse transcription. Taken together, our mapping data reveals the widespread presence of D modification sites on human tRNAs and demonstrates that NaBH_4_-dependent mutational profiling can be utilized for identification and relative measurements of D modification levels.

### Analysis of tRNA expression and codon-specific translation in HEK293T DUS KO cells

To better understand the molecular consequences of D modification on tRNA function and protein translation, we investigated tRNA expression levels and codon-specific translation elongation in DUS1L KO, DUS2L KO, and DUS3L KO cells (HEK293T parent line). All three KO cell lines exhibited reduced cell viability and deficiencies in bulk protein translation (Fig. 3d). First, we generated tRNA sequencing libraries from HEK293T DUS1L, DUS2L, and DUS3L KO cell lines and compared against HEK293T WT cells. Two independent clones (Supplementary. Fig. 3, 4) were analyzed for each DUS KO. Reads were aligned against a custom non-redundant database of mature human tRNA sequences^20^ and DESeq2 was used to measure changes in tRNA expression between WT and KO samples. We clustered tRNA isodecoder data according to common anticodon sequence and correlated statistically significant changes found across both independent KO clones (Fig. 6, Supplementary Datafile 4). Analysis of tRNA expression in DUS1L and DUS3L KO cells indicated only minor defects in tRNA levels. In the DUS1L KO clones, we observed ∼3-fold downregulation of tRNA-Arg-CCT (Fig. 6a, 6b, 6c). According to our NaBH_4_ mapping results tRNA-Arg-CCT contains D at position 17, a likely DUS1L substrate (Fig. 5c); the ∼1000-fold change in mutation rate at this position upon NaBH_4_ treatment was the largest that we measured for any tRNA position. No other tRNA isodecoder families showed consistent downregulation across both DUS1L clones. In DUS3L KO cells, we did not find any tRNA isodecoder families that were downregulated across both KO clones (Fig. 6g, 6h, 6i), although tRNA-Glu-TTC and tRNA-Gly-GCC both showed ∼2.5-fold increase in abundance; neither tRNA contains U at position 47, the major DUS3L substrate site. In contrast to the DUS1L and DUS3L KO cells, we observed stronger downregulation of tRNA levels in DUS2L KO clones, suggesting a more significant role for DUS2L in tRNA stability (Fig. 6d, 6e, 6f). In particular, tRNA-Ser-GCT, tRNA-Pro-AGG, tRNA-Asn-GTT, tRNA-SeC-TCA, and tRNA-Leu-CAG exhibited between ∼4-fold to 8-fold reduction compared to WT cells; all 5 of these tRNAs contain U at position 20, the likely DUS2L substrate site, however we did not detect D modifications in tRNA-Ser-GCT and tRNA-SeC-TCA at this position by NaBH_4_ mapping (Fig. 5c). Some tRNAs were also upregulated in DUS2L KO cells, similar to our findings in DUS1L and DUS3L KO clones. Taken together, our tRNA expression analysis indicates alterations in specific tRNA levels for DUS1L, DUS2L, and DUS3L KO, however only a small group of D-modified tRNAs are affected.

**Figure 6.**
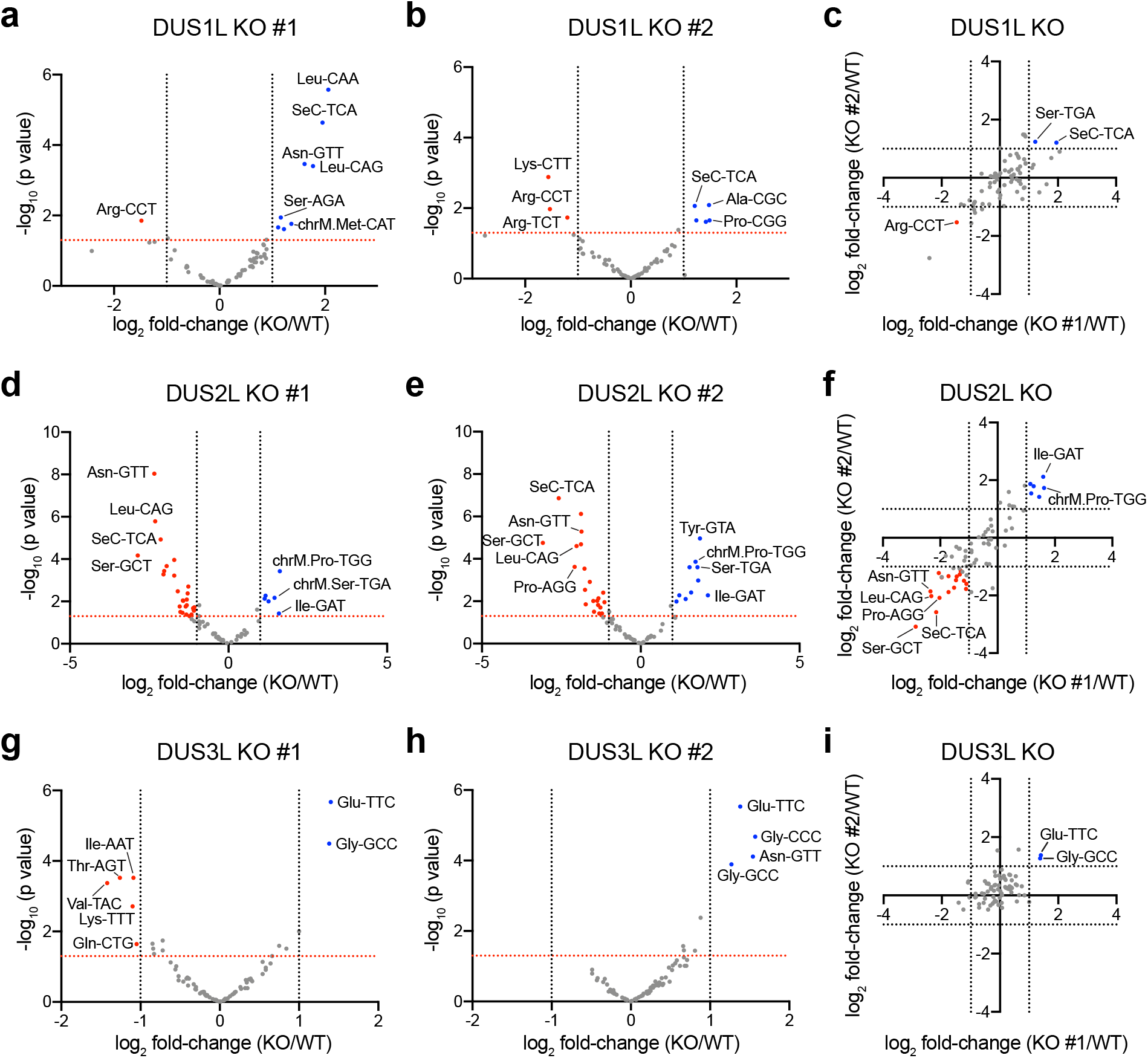
tRNA expression analysis in HEK293T DUS KO cells. **(a)-(b)** Volcano plots of HEK293T DUS1L KO tRNA expression levels across two independent clones. tRNAs with p-values<0.05 and >2-fold expression change are annotated. **(c)** Scatter plot of differentially regulated tRNAs in DUS1L KO clones. tRNAs with >2-fold expression change across both KO clones are annotated. **(d)-(e)** Volcano plots of HEK293T DUS2L KO tRNA expression levels across two independent clones. tRNAs with p-values<0.05 and >2-fold expression change are annotated. **(f)** Scatter plot of differentially regulated tRNAs in DUS2L KO clones. tRNAs with >2-fold expression change across both KO clones are annotated. **(g)-(h)** Volcano plots of HEK293T DUS3L KO tRNA expression levels across two independent clones. tRNAs with p-values<0.05 and >2-fold expression change are annotated. **(i)** Scatter plot of differentially regulated tRNAs in DUS3L KO clones. tRNAs with >2-fold expression change across both KO clones are annotated.

We next investigated codon-specific translation in DUS KO cell lines using a dual luciferase reporter system^20, 41^ containing N*-*terminal *Renilla* luciferase linked upstream to C-terminal firefly luciferase by a linker containing 15 tandem repeats of the same codon (Figure 7a). We picked a variety of codons to assay including cognates for D-modified tRNAs found in our mapping data and tRNAs that showed differences in expression levels in the KO cells. Despite cell proliferation and global translation defects across all 3 DUS KO cell lines, we observed no differences in translation efficiency across most codons that were assayed (Fig. 7b, Supplementary Fig. 16), even for cognate codons of tRNAs that showed decreased expression levels in DUS KOs such as AGC (Ser), AGG (Arg), and AAC (Asn) (Fig. 7b). Only GTT (Val) showed modest reductions (∼10%) in translation efficiency. Interestingly, translation through the related GTG (Val) codon was unaffected in DUS KO cells – while GTG and GTT are decoded by different Val isoacceptors, tRNA-Val-CAC and tRNA-Val-AAC are nearly identical in sequence outside of the anticodon and both contain D modifications (Fig. 5c).

**Figure 7.**
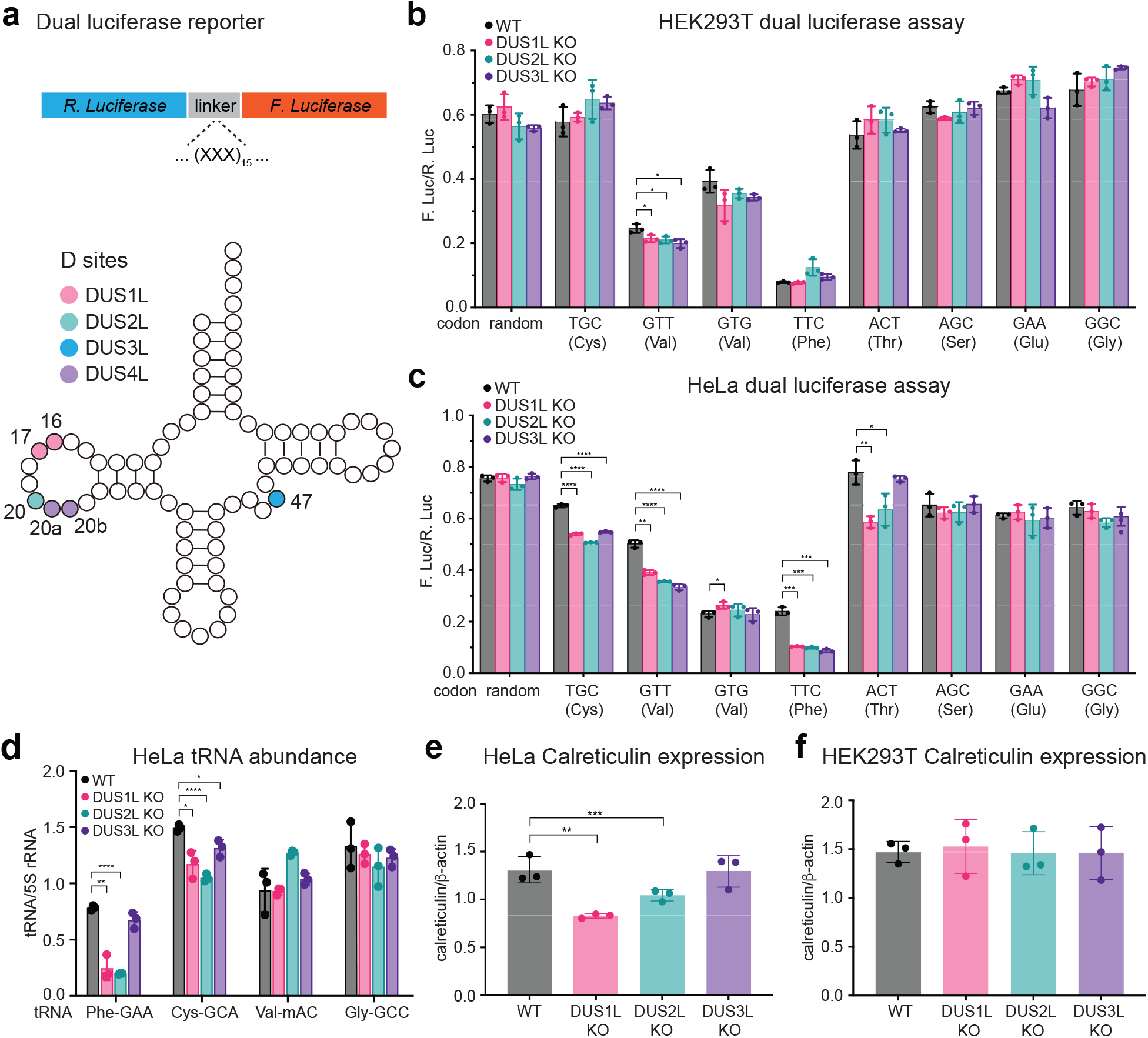
Regulation of codon-dependent translational efficiency by DUS proteins. **(a)** Dual luciferase reporter construct and tRNA schematic showing D sites and associated DUS enzymes. **(b)** Codon-dependent translational efficiency in HEK293T and HEK293T DUS KO cells. Ratio of Firefly luciferase and *Renilla* luciferase signal is shown. Data represent mean values ± s.e.m. Unpaired t-test (two-tailed) were used to evaluate statistical significance. Three independent biological replicates were analyzed per cell line. *P -*values: TTC (Val) WT vs. DUS1L KO, 0.037; TTC (Val) WT vs. DUS2L KO, 0.024; TTC (Val) WT vs. DUS3L KO, 0.014. **(c)** Codon-dependent translational efficiency in HeLa and HeLa DUS KO cells. Ratio of Firefly luciferase and *Renilla* luciferase signal is shown. Data represent mean values ± s.e.m. Unpaired t-test (two-tailed) were used to evaluate statistical significance. Three independent biological replicates were analyzed per cell line. *P-*values: TGC (Cys) WT vs. DUS1L KO, 0.000020; TGC (Cys) WT vs. DUS2L KO, 0.000004; TGC (Cys) WT vs. DUS3L KO, 0.000022; GTT (Val) WT vs. DUS1L KO, 0.00035; GTT (Val) WT vs. DUS2L KO, 0.000064; GTT (Val) WT vs. DUS3L KO, 0.000098; GTG (Val) WT vs. DUS1L, 0.035; TTC (Phe) WT vs. DUS1L, 0.00012; TTC (Phe) WT vs. DUS2L, 0.00011; TTC (Phe) WT vs. DUS2L, 0.00010; ACT (Thr) WT vs. DUS1L, 0.0030; ACT (Thr) WT vs. DUS2L, 0.033. **(d)** Northern blot analysis of tRNA levels in HeLa cells and HeLa DUS KO cells. Data represent mean values ± s.e.m. Unpaired t-test (two-tailed) were used to evaluate statistical significance. Three independent biological replicates were analyzed per cell line. *P*-values: Phe-GAA WT vs. DUS1L, 0.0010; Phe-GAA WT vs. DUS2L, 0.0000016; Cys-GCA WT vs. DUS1L 0.010; Cys-GCA WT vs. DUS2L, 0.000062; Cys-GCA WT vs. DUS3L, 0.017. **(e)** Calreticulin expression in HeLa cells and HeLa DUS KO cell lines. Data represent mean values ± s.e.m. Unpaired t-test (two-tailed) were used to evaluate statistical significance. Three independent biological replicates were analyzed per cell line. *P*-values: WT vs. DUS1L, 0.016; WT vs. DUS2L, 0.00029. **(f)** Calreticulin expression in HEK293T cells and HEK293T DUS KO cell lines. Data represent mean values ± s.e.m. Unpaired t-test (two-tailed) were used to evaluate statistical significance. Three independent biological replicates were analyzed per cell line.

### Knockout of DUS enzymes in HeLa cells

It is known that tRNA pools vary across different human cell lines and tissues^42-45^, and therefore we chose to continue our investigation of D and DUS proteins in HeLa cells. We generated HeLa cell lines containing KOs in DUS1L, DUS2L, or DUS3L (Supplementary Fig. 16, 17) using CRISPR/Cas9, and measured D levels in total RNA and small RNA by LC-QQQ-MS (Supplementary Fig. 18a, 18b, Supplementary Table 7, 8). As in HEK293T cells, D levels were reduced in the individual KO cell lines, although we observed a more dramatic decrease in HeLa for DUS1L as compared to the other KO cell lines and as compared to the HEK293T DUS1L KO (Fig. 3a, 3b). Next, we characterized the proliferation and global protein translation rate in the HeLa DUS KO cell lines. Cell proliferation was decreased in HeLa DUS1L and DUS2L KO cell lines, however HeLa DUS3L KO cells grew at the same rate as WT HeLa (Supplementary. Fig. 18c, Supplementary Table 9). Global protein translation was decreased in all three cell lines (Supplementary Fig. 18d). We also performed NaBH_4_-based mutational profiling of total tRNA extracted from HeLa cells and found ∼50 D sites exhibiting 3-fold or greater NaBH_4_-dependent increases in mutation rate (Supplementary Fig. 19, 20, Supplementary Datafile 5). Overall, the distribution and abundance of D sites in HeLa tRNA closely mirrored our analysis from HEK293T tRNA.

### Codon-specific translation defects in HeLa DUS KO cells

We investigated codon-specific translation in the HeLa DUS KO cell lines using the same panel of dual luciferase reporter constructs as assayed above. In contrast to our results in HEK293T cells, we observed translation defects on multiple codons including GTT (Val), TGC (Cys), ACT (Thr), and TTC (Phe) in DUS KO cells (Fig. 7c, Supplementary Fig. 21). In particular, translation across the 15x TTC (Phe) linker was reduced by ∼50% in DUS1L, DUS2L, and DUS3L KO cell lines as compared to WT HeLa – the most significant reduction in translation of any of the codons assayed. Defects in translation of TGC (Cys) and GTT (Val) were more modest, ranging between 20-30%, but were observed in all three HeLa DUS KO cell lines. Translation through ACT (Thr) was reduced by 20-25% in DUS1L and DUS2L KO lines, but not in DUS3L KO. We were not able to observe translation of the dual luciferase construct containing a 15x TTA (Leu) linker in HeLa cells (Supplementary Fig. 21a) likely due to the low abundance of the corresponding tRNA. Shortening the linker to 5x TTA repeats showed a small (∼12, 15, and 10%), but statistically significant, decrease in translation in HeLa DUS1L, DUS2L, and DUS3L KO (Supplementary Fig. 21b).

To understand whether translation defects in HeLa DUS KO cells were related to alterations in tRNA levels, we extracted total RNA and performed northern blot on tRNAs of interest (Fig. 7d, Supplementary Fig. 22, Supplementary Table 10). We found ∼60% decrease in the abundance of tRNA-Phe-GAA in DUS1L and DUS2L KO HeLa cell lines, but little effect on tRNA-Phe-GAA levels due to DUS3L KO. Interestingly, tRNA-Phe-GAA has U residues at position 16, 17, and 47, but lacks U at position 20, the likely site of DUS2L modification, suggesting a non-enzymatic role for DUS2L in regulating tRNA-Phe-GAA stability. We also measured reductions in levels of tRNA-Cys-GCA across DUS1L, DUS2L, and DUS3L KO cells (Fig. 7d). Levels of tRNA-Val-mAC were unchanged in DUS KO cells. While we observed defects in translation of GTT (Val) codons, sequence similarity among tRNA-Val isoacceptors prevents analysis of specific anticodon species by northern blot. Further, we analyzed the levels of tRNA-Gly-GCC, a tRNA species that showed no translation defects on its cognate codon and did not observe changes in tRNA levels. We also did not observe significant changes in tRNA abundance in HEK293T DUS KO cells for these same tRNAs (Supplementary Fig. 23), consistent with the lack of measurable translation defects in the dual luciferase assay.

After observing translation defects in the dual luciferase reporter assay and decreased abundance of tRNA-Phe-GAA and tRNA-Cys-GCA in HeLa DUS KO cell lines, we hypothesized that genes enriched in these codons may have poorer expression in DUS KO cells compared to WT. Therefore, we analyzed the levels of calreticulin, a calcium-based chaperone protein localized to the endoplasmic reticulum^46^ enriched in Phe residues compared to the proteome average. Western blot of calreticulin in WT and DUS KO HeLa cells demonstrated ∼40% and ∼50% decreased protein levels in HeLa DUS1L and DUS2L KO cell lines, respectively, as compared to WT HeLa, whereas there was no decrease in the DUS3L KO (Fig. 7e, Supplementary Fig. 24). We observed no change in calreticulin levels in the HEK293T DUS KO cell lines (Fig. 7f, Supplementary Fig. 24), consistent with unperturbed translation of TTC (Phe) codons in the dual luciferase assay. Taken together, our work demonstrates that DUS proteins play an important role in translation and tRNA stabilization but indicate that these pathways are dependent on cell type and tRNA identity.

## DISCUSSION

In this manuscript, we investigate the substrates and cellular functions of human DUS proteins. We develop the RNABPP-PS approach, which we apply to show that 5-ClUrd is a general mechanism-based probe for DUS enzymes, and map D sites using catalytic crosslinking and NaBH_4_-based mutational sequencing. We also investigate the role of human DUS proteins in translational regulation and tRNA stability. Our work provides powerful methods for studying DUS proteins and mapping D sites in diverse biological contexts and shows how a ubiquitous tRNA modification functions in a context-specific manner to impact translational efficiency and gene expression.

DUS enzymes and tRNA D modifications are broadly conserved throughout evolution, pointing to an important function in biological systems, but DUS KOs in *E. coli*^10^ and yeast^12^ do not show compromised growth and the effect of D on tRNA function is poorly understood. In contrast, this study and previous work^19^ from our group and others^15, 47^ has shown that human DUS proteins play important roles in cellular proliferation and protein translation. Modifications in the tRNA body often have roles in tRNA stability and folding, which can phenotypically manifest in more subtle ways than anticodon RNA modifications. Whereas our mapping data shows that D is abundant across many tRNAs, only a subset of tRNA species show impaired expression levels upon DUS KO and corresponding translational defects on their cognate codons. In particular, we find codon-specific translational defects at TTC (Phe), TGC (Cys), GTT (Val), and ACT (Thr) codons in HeLa cells, and corresponding decreases in tRNA expression levels for tRNA-Phe-GAA and tRNA-Cys-GCA, suggesting that translation defects result from limiting tRNA levels. Surprisingly, translation of these same codons in HEK293T cells and associated tRNA levels are largely unaffected by DUS KO, even though D modification patterns on tRNAs are largely identical across the two cell lines. Why are only some tRNAs sensitive to loss of D? One explanation is that tRNAs with the highest D modification stoichiometry are most affected by its loss. In addition, differences in tRNA metabolic pathways and tRNA abundance may underly tRNA-specific and cell type-specific sensitivity to DUS KO. In yeast, tRNA hypomodification can trigger tRNA decay pathways^48, 49^, but analogous processes in mammals have not been well studied. Characterizing tRNA surveillance systems in mammals and their interactions with tRNA modifications will be critical to better understand how tRNA levels are regulated. Further, cell type-specific liabilities on individual tRNA species may provide therapeutic opportunities for targeting translation in disease contexts.

Not all of the effects that we observe on tRNA expression levels and codon-specific translation can be rationalized through DUS-mediated D modification. For example, tRNA-Phe-GAA expression in HeLa cells is decreased upon DUS2L KO, however this tRNA lacks a U residue at position 20, the likely site of DUS2L modification. Similarly, position 20 in tRNA-Cys-GCA has been shown to exist as acp^3^U^40^, and our NaBH_4_ mapping supports this modification rather than D; nevertheless, tRNA-Cys-GCA function is sensitive to DUS2L KO in HeLa cells as well. Our data, therefore, suggest that DUS2L may possess some non-catalytic roles and serve as a tRNA chaperone, as has been demonstrated for other tRNA modifying enzymes^50^.

Finally, the RNABPP-PS method, which can be applied on small input quantities and does not require polyadenylated RNA, opens the door to proteomic investigation of RNA modifying enzymes across multiple biological sample types including diverse prokaryotes and mammalian tissues. Combined with a diverse set of modified nucleoside probes^51^ compatible with metabolic labeling, we envision that this strategy will enable new investigations into epitranscriptomic regulatory pathways.

## MATERIALS AND METHODS

### Chemicals

5-FCyd, 5-FUrd, 5-ClCyd, 5-ClUrd, 5-BrCyd, and 5-BrUrd were purchased from Carbosynth. All other chemicals were purchased from Sigma-Aldrich or Fisher Scientific unless noted.

### Plasmids

DUS1L and DUS2L cDNA were obtained from Genscript (NM_022156.5 and NM_017803.5). For construction of Flp-In cell lines, DUS1L and DUS2L cDNA were cloned into a modified pcDNA5/FRT/TO (Life Technologies) vector with an N-terminal 3xFLAG tag. To generate KO cell lines, DNA oligos containing guide RNA (gRNA) sequences were cloned into pX330-U6-Chimeric_BB-CBh-hSpCas9 (Addgene, #42230). The oligos were phosphorylated with T4 PNK (NEB) and annealed and ligated into a BbsI (NEB)-digested pX330 backbone with T4 DNA ligase (NEB). For codon-specific translation experiments, reporter plasmids were generated by replacing the IRES sequence between *Renilla* and Firefly luciferase genes in pCDNA3 RLUC POLIRES FLUC (Addgene #45642) with appropriate 15-codon linkers.

### General cell culture

HEK293T WT and KO cells and HeLa WT and KO cells were cultured at 37°C in a humidified atmosphere with 5% CO2 in DMEM (Thermo Fisher, 11995073) supplemented with 10% fetal bovine serum (Bio-Techne, S1245OH), 1x penicillin-streptomycin (Thermo Fisher, 15070-063) and 2 mM L-glutamine (Thermo Fisher, 25030-081).

### Generation of stable cell lines

Flp-In T-Rex 293 cells were seeded at 0.6 million cells per well in six-well plates and co-transfected with pOG44 (2 µg, Thermo Fisher) and pcDNA5/FRT/TO plasmids (0.2 µg) containing 3xFLAG-DUS1L or 3xFLAG-DUS2L inserts. Cells were selected in 100 µg mL^-1^ hygromycin B and 15 µg mL^-1^ blasticidin and surviving colonies were expanded. To test expression efficiency, cells were induced with tetracycline (0 to 1 µg mL^-1^) for 24 h. Cells were collected and lysed in cell extraction buffer (Invitrogen, FNN0011) with fresh 1 mM phenylmethyl sulfonyl fluoride (PMSF) and protease inhibitor tablets (Sigma, 11836170001). Proteins were separated on SDS-PAGE gels and analyzed by western blotting (anti-FLAG M2, 1:1,000 dilution; Sigma, F1804)

### Generation of KO cell lines

HEK293T WT or HeLa WT cells were seeded at 0.8 million cells per well in six-well plates one day before transfection. 2 µg of pX330 plasmid containing sgRNA for the target protein and 0.2 µg of pcDNA3-FKBP-eGFP-HOTag3 (Addgene, #106924) were co-transfected using Lipofectamine 2000 (Thermo Scientific, 11668027). Cells were sorted by FACS 2 d after transfection. The top 95% of cells displaying GFP signal were sorted as single cells into 96-well dishes. Genomic PCR and western blot (anti-DUS1L, Proteintech, 1:1000; anti-DUS2L, Abcam, 1:1000) were performed to confirm KO.

### Antibodies for western blot

Western blot antibodies: anti-DUS1L, Proteintech, 1:1000; anti-DUS2L, Abcam, 1:1000; anti-calreticulin, Proteintech, 1:1000; anti-FLAG M2, Sigma, 1:1000.

### RNABPP-PS

HEK293T cells were treated with 10 µM 5-FCyd or 250 µM 5-ClCyd at 70% confluency for 16 h. Each replicate used 1 x 10 cm plate. Phase separation-based extraction was performed following literature precedent, with slight modifications^22^. 1 mL of TRIzol reagent (Thermo Fisher, 15596018) was used for lysis per sample, followed by addition of 200 µL of CHCl3. Samples were lysed, vortexed, and centrifuged based on the manufacturer’s protocol. After the first centrifugation, the interphase from each sample was isolated by discarding the aqueous upper phase and the organic lower phase. The interphase was then resuspended in 1 mL of TRIzol for a second isolation step and repeated for three total interphase isolations. After the third extraction, the interphase was resuspended in 100 µL of 100 mM TEAB, 1 mM MgCl2, and 1% SDS buffer and digested with 2 µL RNAse Cocktail (Thermo Fisher, AM2286) for 4 h at 37 °C. An additional 2 µL RNase Cocktail was added overnight to fully digest RNA from crosslinked proteins. The protein concentration in the samples was measured by BCA assay (Thermo Scientific, 23225) and equal amounts of protein were separated on a 10% SDS-PAGE gel followed by western blot analysis. For mass spectrometry analysis, samples were prepared as above but with an additional extraction of TRIzol and RNase Cocktail digestion. 100 µL resuspended sample after the third interphase isolation was subjected to 400 µL of TRIzol and 80 µL of CHCl3. The organic phase was then collected and precipitated using 9 equivalent volumes (900 µL) of methanol. Proteins were resuspended in 100 mM TEAB for proteomic analysis at a final concentration of 1.2 µg µL^-1^.

### Proteomic sample preparation and TMT labeling

Proteomes in 100 mM TEAB buffer (ThermoFisher Scientific) were digested with Lys-C (Wako, 121-05063) for 4 h at RT. The samples were further digested with trypsin (Promega, VA9000) overnight at 37 ^°^C. Digested peptides were labelled with TMTPro reagents (ThermoFIsher Scientific, A44520) for 2 h at RT. Labeling reactions were quenched with 0.5% hydroxylamine for 15 min at RT and subsequently acidified with 0.1% trifluoroacetic acid. Samples were pooled and desalted by C18 stage-tip^52^. Labeled peptides were eluted in 70% acetonitrile, 1% formic acid and dried to completion. Dried peptides were resuspended in 5% acetonitrile, 5% formic acid for LC-MS/MS analysis.

### Proteomic LC-MS/MS

Mass spectrometric data were collected on an Orbitrap Eclipse mass spectrometer (with a FAIMS device enabled) coupled to a Proxeon NanoLC-1000 UHPLC (Thermo Fisher Scientific). The 100 µm capillary column was packed in-house with 35 cm of Accucore 150 resin (2.6 μm, 150Å; ThermoFisher Scientific). Data were acquired for 180 min per run. The scan sequence began with an MS1 scan collected in the orbitrap (resolution: 60,000; scan range: 400−1600 th; automatic gain control (AGC) target: 10e^5^; normalized AGC target: 250%; maximum injection time: 60 ms). MS2 scans were collected in the orbitrap after higher-energy collision dissociation (resolution: 50,000; AGC target: 2e^5^; normalized AGC target: 400%, normalized collision energy (NCE): 37.5; isolation window: 0.5 Th; maximum injection time: 150 ms; top speed set at 1 sec). FAIMS compensation voltages (CVs) were set at -40V, -60V, and -80V.

### Proteomic LC-MS/MS data analysis

Database searching included all entries from the human UniProt Database (downloaded in June 2014). The database was concatenated with one composed of all protein sequences for that database in the reversed order^53^. Raw files were converted to mzXML, and monoisotopic peaks were reassigned using Monocle^54^. Searches were performed with Comet^55^ using a 50 ppm precursor ion tolerance and fragment bin tolerance of 0.02. TMTpro labels on lysine residues and peptide N-termini (+304.207 Da) and oxidation of methionine residues (+15.995 Da) were set as variable modifications. Peptide-spectrum matches (PSMs) were adjusted to a 1% false discovery rate (FDR) using a linear discriminant after which proteins were assembled further to a final protein-level FDR of 1% analysis^56^. TMT reporter ion intensities were measured using a 0.003 Da window around the theoretical *m/z* for each reporter ion. Proteins were quantified by summing reporter ion counts across all matching PSMs. More specifically, reporter ion intensities were adjusted to correct for the isotopic impurities of the different TMTpro reagents according to manufacturer specifications. Peptides were filtered to exclude those with a summed signal-to-noise (SN)<150 across all TMT channels and <0.5 precursor isolation specificity. The signal-to-noise (S/N) measurements of peptides assigned to each protein were summed (for a given protein).

### Nucleoside LC-QQQ-MS

For measurement of the metabolic incorporation of C5-modified halopyrimidine nucleosides, HEK293T WT cells were treated with 10 µM 5-FCyd, 250 µM 5-ClCyd, or 500 µM 5-BrCyd for 16 h overnight, and total RNA was extracted with TRIzol reagent using the manufacturer’s protocol. For measurement of native RNA modification levels in WT and KO cells, total RNA was isolated as above and additionally fractionated into small RNA using RNA Clean & Concentrator-5 (Zymo) following the manufacturer’s protocol. RNA samples were digested and dephosphorylated to nucleosides and analyzed by LC-QQQ-MS on an Agilent 6470 following literature precedent^19^. MS1 (parent ion) to MS2 (deribosylated base ion) transition for modified nucleosides were set as follows: *m/z* 262 → 130 for 5-FCyd, *m/z* 263 → 131 for 5-FUrd; *m/z* 278 → 146 for 5-ClCyd; *m/z* 279 → 147 for 5-ClUrd; *m/z* 322 → 190 for 5-BrCyd; *m/z* 323 → 191 for 5-BrUrd; *m/z* 247 → 115 for D.

### Cell proliferation assay

Cells were plated in a 96-well plate (4000 cells in 200 µL media per well) on day 0. Cell proliferation was measured daily using the MTS assay (CellTiter 96 Aqueous Non-Radioactive Cell Proliferation Assay; Promega, G5430) across 3 d (days 1 – 3). Absorbance was read at 490 nm with a Spectramax iD5 (Molecular Devices).

### Global translation assay

Global translation efficiency was assayed using O-propargyl puromycin (OP puro, Click Chemistry Tools) incorporation and measuring fluorescence after click chemistry via flow cytometry^19, 57^ or fluorescence microscopy^19, 58^ following literature precedent. For fluorescence microscopy quantification of OP-puro signal, we used Fiji ImageJ (2.14.0/1.54f). Cells masks were generated using DAPI staining and overlaid with the Cy3 signal to measure intensity in individual cells. The resulting measurement of mean intensity density over area was used as comparison of translational efficiency across cell lines.

### 5-ClUrd-iCLIP and bioinformatic analysis

5-ClUrd-iCLIP for DUS1L and DUS2L was performed following literature precedent^19, 59^. Flp-In T-Rex 293 cells expressing 3xFLAG-DUS1L or 3xFLAG-DUS2L were treated with 100 μM 5-ClUrd in regular growth medium for 12 h. Untreated cells expressing DUS1L or DUS2L were used as the control. 5-ClUrd-iCLIP data was processed following literature precedent^19^ using the iCount Primary Analysis pipeline (consensus mapping) on the iMAPs web server (https://imaps.genialis.com/iclip). Two independent replicates of 5-ClUrd and control samples were analyzed. Crosslinking site read counts were used for peak calling analysis by Paraclu with the following parameters: a minimal sum of scores inside a cluster of 10, a maximal cluster size of 5, and a minimal density increase of 2. The peaks were further intersected with crosslinking sites. We only kept peaks that overlapped with, or were 1 nt adjacent to, U residues for further analysis. Read counts were normalized per million uniquely mapping reads (RPM). Peaks that were unique to the 5-ClUrd-treated sample or showed fold change >4 as compared to the control were kept after intersection. Only peaks with read counts >10 in two replicates were selected for further analyses. Sequences five bases upstream and downstream were extracted from the reference and used for motif analysis by MEME (V.5.3.3) using -mod zoops - nmotifs 3 -minw 6 -maxw 50 -objfun classic -revcomp -markov_order 0

### tRNA sequencing sample preparation

Total RNA was isolated from HEK293T WT and DUS KO cells by TRIzol extraction according to the manufacturer’s protocol. RNA was separated on a 10% TBE-urea gel and bands corresponding to mature tRNA (∼75 nt) were excised. tRNA was eluted by passive diffusion in TE buffer (pH 7.5) at room temperature overnight and recovered by EtOH precipitation. AlkB treatment was performed to remove common tRNA modifications following literature precedent^37^. Libraries were prepared using the NEBNext® Small RNA Library Prep Set for Illumina® (NEB #E7330S) following the manufacturer’s protocols as described previously^36^. The barcoded PCR products were separated on a 5% native TBE-PAGE gel and bands corresponding to full-length mature tRNA (∼200 nt) were excised and purified. Barcoded samples were pooled and submitted for sequencing on a NovaSeq 6000 (Illumina). To identify D sites via NaBH4-induced mutations, gel purified tRNA was reduced by NaBH4 prior to AlkB treatment: in a 100 μL reaction, 5 µg of tRNA was reacted with NaBH4 (10 mg/mL final concentration, 100 mg/mL stock in 10 mM KOH) in 40 mM Tris•HCl pH 7.5 on ice for 1 h. The reaction was quenched with 750 mM acetic acid and RNA was purified by EtOH precipitation. During the library preparation, reverse transcription was performed with Superscript II reverse transcriptase (Thermo Scientific) at 42 °C. For tRNA expression analysis, in vitro transcribed (IVT) *E. coli* tRNAs were spiked into total RNA for normalization purpose. In brief, 30 ng each of three IVT *E. coli* tRNAs (tRNA-Tyr, tRNA-Phe, and tRNA-Lys)^37^ were added to 100 μg of total RNA (∼0.01 pmol of each tRNA/1 μg of total RNA) and the sample was deacylated by heating at 37 °C for 45 min in 100 mM Tris•HCl pH 9.0 and 1 mM EDTA prior to gel purification.

### Bioinformatic analysis of tRNA sequencing

Demultiplexed tRNA sequencing data was uploaded to Galaxy and Fastp was used to trim adaptors and filter out low-quality reads. The adaptor sequence used for input 1 was: AGATCGGAAGAGCACACGTCTGAACTCCAGTCAC; the adaptor sequence used for input 2 was: GATCGTCGGACTGTAGAACTCTGAACGTGTAGATCTCGGTGGTCGCCGTATCATT. Trimmed reads were mapped with RNA STAR Gapped-read mapper for RNA-seq data (Galaxy Version 2.7.8a+galaxy0) to a custom non-redundant human mature tRNA reference as described previously^20^. For tRNA expression analysis, the three spike-in *E. coli* tRNA sequences were added to the human tRNA reference. The length of the SA pre-indexing string was set as 6; default values were used for all other parameters. To identify D sites via NaBH4-induced mutations, three independent biological samples were sequenced for HEK293T and HEK293T DUS3L KO tRNA, and two independent replicates for HeLa tRNA. BAM files from independent replicates were merged before mutational analysis with VarScan Somatic^60^. The mature tRNA reference was used as the reference genome, untreated samples were used as the input for the “normal sample”, and the NaBH4 treated samples were used as the input for the “tumor sample”. The estimated purity of the normal sample was set to 1, and the estimated purity of the tumor sample was set to 0.1. The minimum length of ref-supporting reads and the minimum length of variant-supporting reads were both set to 15. Default settings were used for all other parameters. Positions with less than 50 total reads and p≥0.05 were discarded. Mutation rates for individual positions within isodecoder families were calculated by summing mutated and unmutated reads from tRNA isotypes with identical anticodon sequence. To focus on potential D sites, a list of putative D sites on tRNA was curated manually. For tRNA expression analysis, uniquely mapped reads for each tRNA isotype were counted using featureCounts. DESeq2 analysis was performed to calculate the fold change of each tRNA in the KO samples versus the WT. The normalization factors of the analysis were set as the three spike-in *E. coli* tRNAs. For analysis at the anticodon level (isodecoder families), read numbers for each anticodon were summed up with Datamash. DESeq2 was performed as described above for the isotype analysis.

### Dual luciferase reporter assay

To measure codon-specific translational efficiencies, cells were plated in a 24-well plate (40,000 cells per well) on day 0. The following day, 600 ng of appropriate reporter plasmid was transfected with Lipofectamine 2000. A reporter plasmid containing a linker with 15 random codons was used as a control. Cells were lysed in passive lysis buffer (Promega) following the manufacturer’s protocol and transferred to a 96-well plate. Firefly and *Renilla* luciferase activity were measured using the Dual-luciferase Reporter Assay (Promega) and luciferase signal was measured with a Spectramax iD5 (Molecular Devices). The ratio of Firefly/*Renilla* luciferase signal was used to evaluate translational efficiency through the linker region of each reporter. Three independent biological replicates were analyzed for each sample.

### tRNA abundance northern blot analysis

Measurements of tRNA abundance were performed by northern blot analysis. In brief, total RNA was extracted using TRIzol, separated on an 8% TBE-urea gel, and transferred via capillary action to Biodyne B Nylon Membrane (Thermo Fisher) overnight. The following day, RNA was crosslinked to the membrane by irradiation with a 254 nm UV lamp for 15 min. The membrane was then blocked with 5 µg sheared salmon sperm DNA in 200 mM Na2HPO4, pH 7.2 with 7% SDS (Thermo Fisher) for 1 h at 45 °C. Next, 50 pmol/mL of a biotinylated tRNA antisense probe and 50 pmol/mL of IR800-labeled 5S RNA antisense probe were hybridized to the blot in 200 mM Na2HPO4, pH 7.2 with 7% SDS for 4 h at 45 °C. The membrane was washed three times with 2X SSC buffer with 0.1% SDS at room temperature. Hybridized biotinylated antisense probe was labeled with IR680 dye in 2X SSC, 0.1% SDS (BioRad) for 1 h at room temperature. The membrane was washed again three times and imaged.

## ACKNOWLEDGMENTS

We thank Christina DeCoste at the Princeton University Flow Cytometry Resource Facility for assistance with FACS analysis, and Wei Wang and Lance Parsons at the Princeton University Genomics Core Facility and Lewis Sigler Institute for assistance with high-throughput sequencing and bioinformatic analysis. We thank Venu Vandavasi at the Princeton University Biophysics Core facility for assistance with plate reader measurements. We thank Rachel Rodrigues at the Thermo Fisher Scientific Center for Multiplexed Proteomics at Harvard Medical School (https://tcmp.hms.harvard.edu) for performing proteomics analysis of RNABPP-PS samples. R.E.K. acknowledges support from the NIH (R01 GM132189), NSF (MCB-1942565), Alfred P. Sloan Foundation, and the Princeton Catalysis Initiative. N.J.Y. and W.D. were supported by a generous gift from the Edward C. Taylor 3rd Year Graduate Fellowship in Chemistry. All authors acknowledge financial support from Princeton University.

