## Supplementary Information for "Cell type-specific translational regulation by human DUS enzymes"

Table of contents

Supplementary Tables 1-10 pg 2

Supplementary Figures 1-24 pg 12

**Supplementary Table 1.** LC-QQQ-MS analysis of 5-halo-pyrimidine labeling of total RNA in HEK293T WT cells. Cells were treated with 10 μM 5-FCyd, 250 μM 5-ClCyd, or 500 μM 5-BrCyd for 16 h overnight.

| treatment condition | nucleoside concentration (ng/mL) | | | | | | 5-XCyd /C % | 5-XUrd /C % |
|---|---|---|---|---|---|---|---|---|
| | A | G | C | U | 5-XCyd | 5-XUrd | | |
| 10 μM 5-FCyd_1 | 46509.32 | 73614.74 | 70408.98 | 37312.31 | 412.77 | 267.34 | 0.59 | 0.38 |
| 10 μM 5-FCyd_2 | 21472.80 | 36054.27 | 30596.33 | 16752.15 | 168.41 | 124.71 | 0.55 | 0.41 |
| 10 μM 5-FCyd_3 | 10188.55 | 19609.63 | 14444.55 | 7114.84 | 100.30 | 64.83 | 0.69 | 0.45 |
| 250 μM 5-ClCyd_1 | 37932.63 | 76884.01 | 56497.74 | 32347.08 | 238.16 | 243.55 | 0.42 | 0.43 |
| 250 μM 5-ClCyd_2 | 40264.86 | 80610.37 | 60996.06 | 34176.63 | 245.92 | 280.62 | 0.40 | 0.46 |
| 250 μM 5-ClCyd_3 | 51177.79 | 101579.54 | 80196.99 | 44832.08 | 259.36 | 308.78 | 0.32 | 0.39 |
| 500 μM 5-BrCyd_1 | 15471.37 | 30121.35 | 22907.84 | 11820.09 | 255.26 | 172.89 | 1.11 | 0.75 |
| 500 μM 5-BrCyd_2 | 30151.49 | 53430.01 | 46383.11 | 23759.79 | 489.91 | 299.94 | 1.06 | 0.65 |
| 500 μM 5-BrCyd_3 | 18244.92 | 33849.62 | 24779.82 | 13197.91 | 303.43 | 189.98 | 1.22 | 0.77 |

**Supplementary Table 2.** Guide RNA sequences for CRISPR-Cas9 genome editing.

| | |
|---|---|
| **DUS1L** | CCTTCCGGTAGTTGGCGTCG |
| **DUS2L** | GAAGACAACTCGATCATCAG (sgRNA 1)<br>TAACACACCATCCTTACCGA (sgRNA 2) |
| **DUS3L** | GACTTTGTGGACATCAACGT |
| **DUS4L** | GCAATGGTTCGATATTCAAAG |

**Supplementary Table 3.** LC-QQQ-MS analysis of D levels in total RNA in HEK293T cells.

| | nucleoside concentration (ng/mL) | | | | | D/U % |
|---|---|---|---|---|---|---|
| | A | G | C | U | D | |
| WT_1 | 18700.72 | 36000.46 | 29533.87 | 1708.50 | 17.85 | 1.04 |
| WT_2 | 15016.65 | 29298.68 | 23675.37 | 1498.57 | 15.13 | 1.01 |
| WT_3 | 13107.71 | 24695.73 | 21590.02 | 1233.43 | 13.24 | 1.07 |
| DUS1L KO _1 | 8348.79 | 15773.88 | 14472.04 | 771.87 | 5.25 | 0.68 |
| DUS1L KO _2 | 14701.30 | 27107.39 | 24791.07 | 1363.74 | 9.65 | 0.71 |
| DUS1L KO _3 | 13736.46 | 25322.71 | 23528.79 | 1312.03 | 9.19 | 0.70 |
| DUS2L KO_1 | 11569.92 | 21283.89 | 20071.59 | 1113.62 | 7.38 | 0.66 |
| DUS2L KO_2 | 14306.43 | 26216.30 | 23371.37 | 1374.68 | 8.44 | 0.61 |
| DUS2L KO_3 | 11787.31 | 22208.34 | 20350.93 | 1171.04 | 7.49 | 0.64 |
| DUS3L KO_1 | 12608.22 | 24694.41 | 22013.33 | 1190.46 | 9.80 | 0.82 |
| DUS3L KO_2 | 15594.58 | 30154.56 | 26929.49 | 1483.78 | 12.59 | 0.85 |
| DUS3L KO_3 | 15610.97 | 29970.54 | 27657.42 | 1520.65 | 13.31 | 0.88 |
| DUS1L + DUS2L KO_1 | 10677.03 | 19565.99 | 19035.06 | 1035.24 | 3.73 | 0.36 |
| DUS1L + DUS2L KO _2 | 16213.63 | 29638.20 | 26347.33 | 1452.35 | 5.92 | 0.41 |
| DUS1L + DUS2L KO _3 | 15939.41 | 28894.03 | 27276.26 | 1523.07 | 5.77 | 0.38 |
| DUS quadruple KO_1 | 16067.77 | 29893.94 | 28064.21 | 1517.73 | 0.00 | 0.00 |
| DUS quadruple KO_2 | 17903.63 | 33321.31 | 31345.93 | 1718.92 | 0.00 | 0.00 |
| DUS quadruple KO_3 | 15903.76 | 29437.51 | 29316.05 | 1542.51 | 0.00 | 0.00 |

**Supplementary Table 4.** LC-QQQ-MS analysis of D levels in small RNA in HEK293T cells

| | nucleoside concentration (ng/mL) | | | | | D/U % |
|---|---|---|---|---|---|---|
| | A | G | C | U | D | |
| WT_1 | 1492.12 | 2423.59 | 669.15 | 979.61 | 104.41 | 10.66 |
| WT_2 | 1326.86 | 2108.87 | 556.79 | 855.61 | 86.25 | 10.08 |
| WT_3 | 1451.88 | 2280.43 | 630.90 | 964.75 | 99.07 | 10.27 |
| DUS1L KO _1 | 806.91 | 1186.31 | 366.04 | 517.86 | 30.48 | 5.89 |
| DUS1L KO _2 | 1264.95 | 1904.16 | 613.80 | 850.55 | 56.10 | 6.60 |
| DUS1L KO _3 | 1322.20 | 1999.38 | 628.89 | 907.55 | 57.71 | 6.36 |
| DUS2L KO_1 | 867.36 | 1233.67 | 368.61 | 550.67 | 30.87 | 5.61 |
| DUS2L KO_2 | 1161.62 | 1701.59 | 535.34 | 797.93 | 44.66 | 5.60 |
| DUS2L KO_3 | 966.88 | 1391.08 | 429.42 | 641.70 | 35.12 | 5.47 |
| DUS3L KO_1 | 1001.36 | 1517.17 | 454.39 | 640.41 | 48.02 | 7.50 |
| DUS3L KO_2 | 1336.20 | 2095.62 | 683.14 | 921.68 | 73.38 | 7.96 |
| DUS3L KO_3 | 1311.59 | 2027.13 | 618.99 | 912.37 | 70.65 | 7.74 |
| DUS1L + DUS2L KO_1 | 1041.58 | 1511.97 | 480.66 | 705.48 | 20.60 | 2.92 |
| DUS1L + DUS2L KO _2 | 1323.07 | 1995.15 | 586.64 | 941.02 | 29.97 | 3.18 |
| DUS1L + DUS2L KO _3 | 1243.42 | 1848.63 | 592.68 | 867.31 | 26.57 | 3.06 |
| DUS quadruple KO_1 | 1711.69 | 2760.88 | 756.61 | 1298.21 | 0.00 | 0.00 |
| DUS quadruple KO_2 | 1446.33 | 2260.27 | 754.36 | 1081.13 | 0.00 | 0.00 |
| DUS quadruple KO_3 | 1404.58 | 2206.60 | 722.55 | 1052.09 | 0.00 | 0.00 |

**Supplementary Table 5.** Cell proliferation data for HEK293T, DUS1L KO, DUS2L KO, DUS1L + DUS2L KO, and ΔDUS KO. Absorbance at 490 nm was measured using the MTS assay and normalized to the average of HEK293T WT values on day 3. Four independent biological replicates were prepared for each cell line, with three technical replicates per sample.

| **HEK 293T WT** | | | **DUS1L KO** | | | **DUS2L KO** | | | **DUS1L + DUS2L KO** | | | **ΔDUS KO** | | |
|---|---|---|---|---|---|---|---|---|---|---|---|---|---|---|
| **day 1** | **day 2** | **day 3** | **day 1** | **day 2** | **day 3** | **day 1** | **day 2** | **day 3** | **day 1** | **day 2** | **day 3** | **day 1** | **day 2** | **day 3** |
| 14.67 | 47.46 | 86.48 | 11.49 | 36.00 | 59.15 | 9.33 | 27.90 | 55.86 | 11.22 | 24.94 | 50.50 | 6.15 | 19.42 | 31.15 |
| 14.62 | 45.95 | 99.93 | 12.44 | 33.32 | 58.65 | 9.55 | 24.83 | 54.93 | 10.79 | 23.41 | 50.83 | 5.58 | 20.77 | 39.80 |
| 15.14 | 45.56 | 87.70 | 11.99 | 33.42 | 60.42 | 9.69 | 24.47 | 54.52 | 11.25 | 24.00 | 53.01 | 6.54 | 17.88 | 37.49 |
| 15.50 | 49.37 | 99.31 | 9.15 | 37.64 | 84.75 | 11.99 | 27.38 | 68.99 | 9.36 | 27.71 | 56.86 | 5.77 | 16.54 | 41.53 |
| 15.09 | 49.94 | 95.89 | 10.55 | 37.23 | 76.81 | 10.64 | 22.56 | 65.90 | 8.57 | 29.70 | 53.37 | 5.96 | 20.00 | 33.65 |
| 15.34 | 49.30 | 102.62 | 9.93 | 37.76 | 76.00 | 10.67 | 22.54 | 66.09 | 9.29 | 31.62 | 54.84 | 5.96 | 21.15 | 39.99 |
| 14.29 | 47.37 | 101.57 | 11.74 | 34.37 | 87.66 | 12.67 | 28.96 | 77.22 | 4.14 | 33.87 | 66.12 | 6.73 | 17.50 | 37.49 |
| 14.77 | 48.76 | 100.65 | 9.50 | 35.13 | 79.79 | 10.55 | 25.40 | 65.84 | 2.74 | 32.54 | 62.04 | 5.96 | 16.73 | 38.26 |
| 14.03 | 50.14 | 101.22 | 8.95 | 35.12 | 77.77 | 10.56 | 25.50 | 66.48 | 2.90 | 33.12 | 61.16 | 6.15 | 20.77 | 33.84 |
| 18.11 | 39.24 | 106.29 | 8.62 | 29.56 | 82.14 | 13.36 | 22.09 | 76.95 | 11.67 | 25.53 | 60.51 | 6.15 | 20.96 | 37.69 |
| 17.17 | 37.86 | 107.23 | 10.64 | 29.93 | 74.64 | 11.88 | 21.80 | 70.43 | 9.23 | 26.99 | 56.86 | 7.11 | 17.88 | 37.69 |
| 17.79 | 38.60 | 111.11 | 10.08 | 30.38 | 74.88 | 11.97 | 21.83 | 71.56 | 9.10 | 27.48 | 59.21 | 6.35 | 17.11 | 38.07 |

**Supplementary Table 6.** Cy3 median values for HEK293T WT, DUS1L KO, and DUS2L KO in OP-puro global protein translation assay. Four independent biological replicates were prepared for each cell line, with three technical replicates per sample. Averages of each biological replicate are shown. 50,000 cells were analyzed per run.

| **Cy3 Median** | | |
|---|---|---|
| **WT** | **DUS1L KO** | **DUS2L KO** |
| 21906.7 | 17404.67 | 18910 |
| 21987.3 | 16321.67 | 19809 |
| 24677.3 | 15517.33 | 17117.33 |
| 24248 | 14348.33 | 19456.67 |

**Supplementary Table 7.** LC-QQQ-MS analysis of D levels in HeLa total RNA

| | nucleoside concentration (ng/mL) | | | | | D/U % |
|---|---|---|---|---|---|---|
| | A | G | C | U | D | |
| WT_1 | 1942.23 | 5001.27 | 4349.36 | 1078.99 | 17.11 | 1.59 |
| WT_2 | 2996.13 | 7217.59 | 6439.37 | 1540.38 | 22.01 | 1.43 |
| WT_3 | 1239.23 | 3159.72 | 2799.35 | 666.81 | 11.67 | 1.75 |
| DUS1L KO _1 | 2732.60 | 6508.36 | 6723.76 | 1426.26 | 8.16 | 0.57 |
| DUS1L KO _2 | 3625.17 | 8354.43 | 7786.46 | 3873.25 | 22.44 | 0.58 |
| DUS1L KO _3 | 2461.82 | 5888.09 | 6024.27 | 3195.20 | 18.25 | 0.57 |
| DUS2L KO_1 | 2157.95 | 5612.42 | 5019.60 | 1184.04 | 12.16 | 1.03 |
| DUS2L KO_2 | 1615.14 | 4073.89 | 3785.44 | 868.35 | 9.91 | 1.14 |
| DUS2L KO_3 | 3524.62 | 8276.90 | 7943.11 | 1823.52 | 16.55 | 0.91 |
| DUS3L KO_1 | 2972.13 | 7279.08 | 7317.03 | 1562.57 | 15.89 | 1.02 |
| DUS3L KO_2 | 3239.71 | 7758.66 | 6924.01 | 1627.73 | 20.76 | 1.28 |
| DUS3L KO_3 | 2746.99 | 6724.89 | 6539.11 | 1424.36 | 16.37 | 1.15 |

**Supplementary Table 8.** LC-QQQ-MS analysis of D levels in HeLa small RNA.

| | nucleoside concentration (ng/mL) | | | | | D/U % |
|---|---|---|---|---|---|---|
| | A | G | C | U | D | |
| WT_1 | 288.60 | 600.98 | 509.23 | 152.28 | 21.68 | 14.24 |
| WT_2 | 386.37 | 861.40 | 663.58 | 207.82 | 29.14 | 14.02 |
| WT_3 | 400.21 | 951.40 | 709.52 | 228.21 | 27.86 | 12.21 |
| DUS1L KO _1 | 392.33 | 537.08 | 619.56 | 169.35 | 6.83 | 4.03 |
| DUS1L KO _2 | 674.71 | 825.17 | 528.01 | 246.71 | 11.62 | 4.71 |
| DUS1L KO _3 | 745.94 | 1577.67 | 1086.85 | 816.84 | 33.20 | 4.06 |
| DUS2L KO_1 | 357.02 | 852.36 | 580.30 | 195.20 | 18.09 | 9.27 |
| DUS2L KO_2 | 726.86 | 1585.64 | 1192.71 | 410.29 | 33.59 | 8.19 |
| DUS2L KO_3 | 424.87 | 978.04 | 827.53 | 240.78 | 20.69 | 8.59 |
| DUS3L KO_1 | 413.52 | 889.50 | 788.19 | 227.09 | 24.38 | 10.73 |
| DUS3L KO_2 | 507.80 | 1124.08 | 1135.00 | 285.31 | 27.60 | 9.68 |
| DUS3L KO_3 | 648.27 | 1379.54 | 1021.79 | 359.44 | 36.15 | 10.06 |

**Supplementary Table 9.** Cell proliferation data for HeLa, DUS1L KO, DUS2L KO, and DUS3L KO. Absorbance at 490 nm was measured using the MTS assay and normalized to the average of HeLa WT values on day 3. Four independent biological replicates were prepared for each cell line, with three technical replicates per sample

| **HeLa WT** | | | **DUS1L KO** | | | **DUS2L KO** | | | **DUS3L KO** | | |
|---|---|---|---|---|---|---|---|---|---|---|---|
| **day 1** | **day 2** | **day 3** | **day 1** | **day 2** | **day 3** | **day 1** | **day 2** | **day 3** | **day 1** | **day 2** | **day 3** |
| 24.82 | 73.19 | 98.45 | 25.56 | 55.88 | 87.38 | 22.71 | 67.59 | 91.95 | 22.74 | 73.52 | 92.41 |
| 23.24 | 74.29 | 91.83 | 22.94 | 56.31 | 68.74 | 24.28 | 74.28 | 100.98 | 27.81 | 70.07 | 113.89 |
| 23.71 | 73.17 | 95.94 | 24.04 | 59.41 | 84.98 | 25.52 | 76.41 | 93.38 | 27.77 | 68.98 | 111.44 |
| 25.06 | 69.47 | 105.95 | 24.62 | 51.43 | 86.20 | 26.77 | 67.05 | 84.67 | 23.47 | 73.60 | 105.08 |
| 24.80 | 72.26 | 100.67 | 25.87 | 56.06 | 92.14 | 23.08 | 54.44 | 80.92 | 21.56 | 69.21 | 93.17 |
| 23.40 | 68.82 | 94.85 | 23.51 | 56.02 | 76.44 | 22.26 | 50.97 | 81.08 | 24.54 | 66.00 | 103.76 |
| 24.12 | 69.22 | 102.26 | 23.84 | 56.70 | 91.00 | 22.97 | 54.29 | 79.86 | 26.07 | 62.74 | 103.90 |
| 25.83 | 71.28 | 105.47 | 24.55 | 50.89 | 86.52 | 23.72 | 57.33 | 81.83 | 21.79 | 66.36 | 101.24 |
| 24.79 | 73.65 | 98.12 | 25.70 | 57.89 | 94.64 | 23.16 | 53.15 | 81.41 | 21.81 | 68.04 | 94.84 |
| 23.48 | 69.88 | 99.31 | 23.74 | 57.53 | 81.03 | 21.75 | 49.11 | 80.46 | 24.35 | 67.13 | 104.55 |
| 24.38 | 70.46 | 101.37 | 23.83 | 57.94 | 91.51 | 22.47 | 53.43 | 78.44 | 26.01 | 63.44 | 104.71 |
| 26.27 | 73.74 | 105.79 | 25.11 | 52.26 | 86.01 | 23.45 | 55.98 | 80.36 | 22.00 | 66.68 | 100.95 |

**Supplementary Table 10.** Sequences of labeled probes for tRNA northern blotting analyses. Bio = biotin

| RNA | Sequence |
|---|---|
| tRNA-Phe-GAA | 5'-Bio-TGGTGCCGAAACCCGGGATTGAACCGGGG |
| tRNA-Cys-GCA | 5'-Bio-AGTCAAATGCTCTACCACTGAGCTATACCCCC |
| tRNA-Val-mAC | 5'-Bio-TGTTTCCGCCCGGTTTCGAACCGGGGACCTTTCGCGT |
| tRNA-Gly-GCC | 5'-Bio-GCAGGCGAGAATTCTACCACTGAACCACCCATGC |

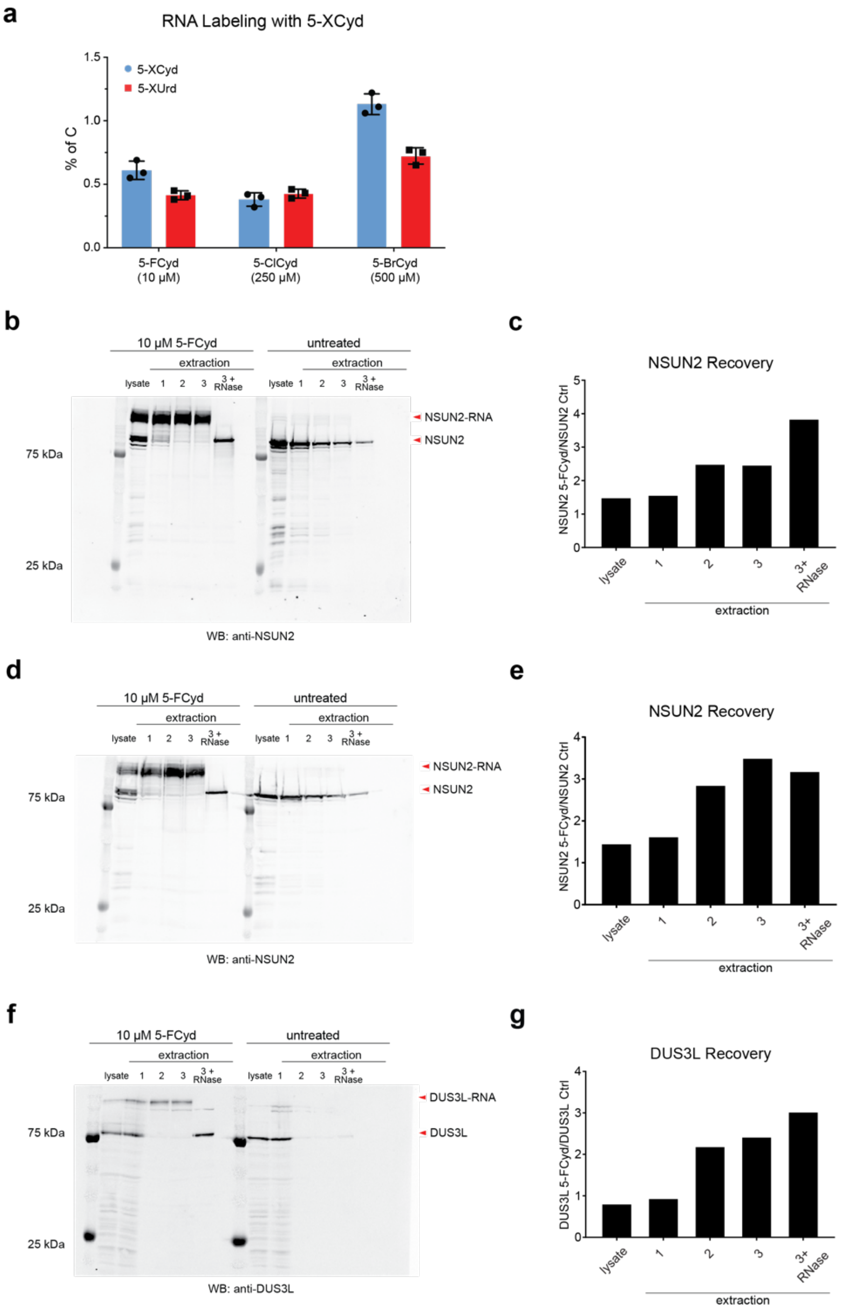


**Supplementary Figure 1.** Validation of RNABPP-PS with 5-FCyd treated cells. **(a)** Metabolic incorporation measurements for 5-FCyd/5-FUrd, 5-ClCyd/5-ClUrd, and 5-BrCyd/5-BrUrd. Raw values are shown in Supplementary Table 1. **(b)** Western blot recovery and enrichment of NSUN2 in 10 µM 5-FCyd-treated HEK293T cells versus control upon sequential interphase isolations. **(c)** Fold change of **(b)** NSUN2 recovery in interphase with 10 µM 5-FCyd treatment versus no treatment. **(d)** Second replicate of western blot recovery and enrichment of NSUN2 in 10 µM 5-FCyd-treated HEK293T cells versus control upon sequential interphase isolations. **(e)** Fold change of **(d)** NSUN2 recovery in interphase with 10 µM 5-FCyd treatment versus no treatment. **(f)** Western blot recovery and enrichment of DUS3L in 10 µM 5-FCyd-treated HEK293T cells versus control upon sequential interphase isolations. **(g)** Fold change of DUS3L recovery in interphase with 250 µM 5-ClCyd treatment versus no treatment.

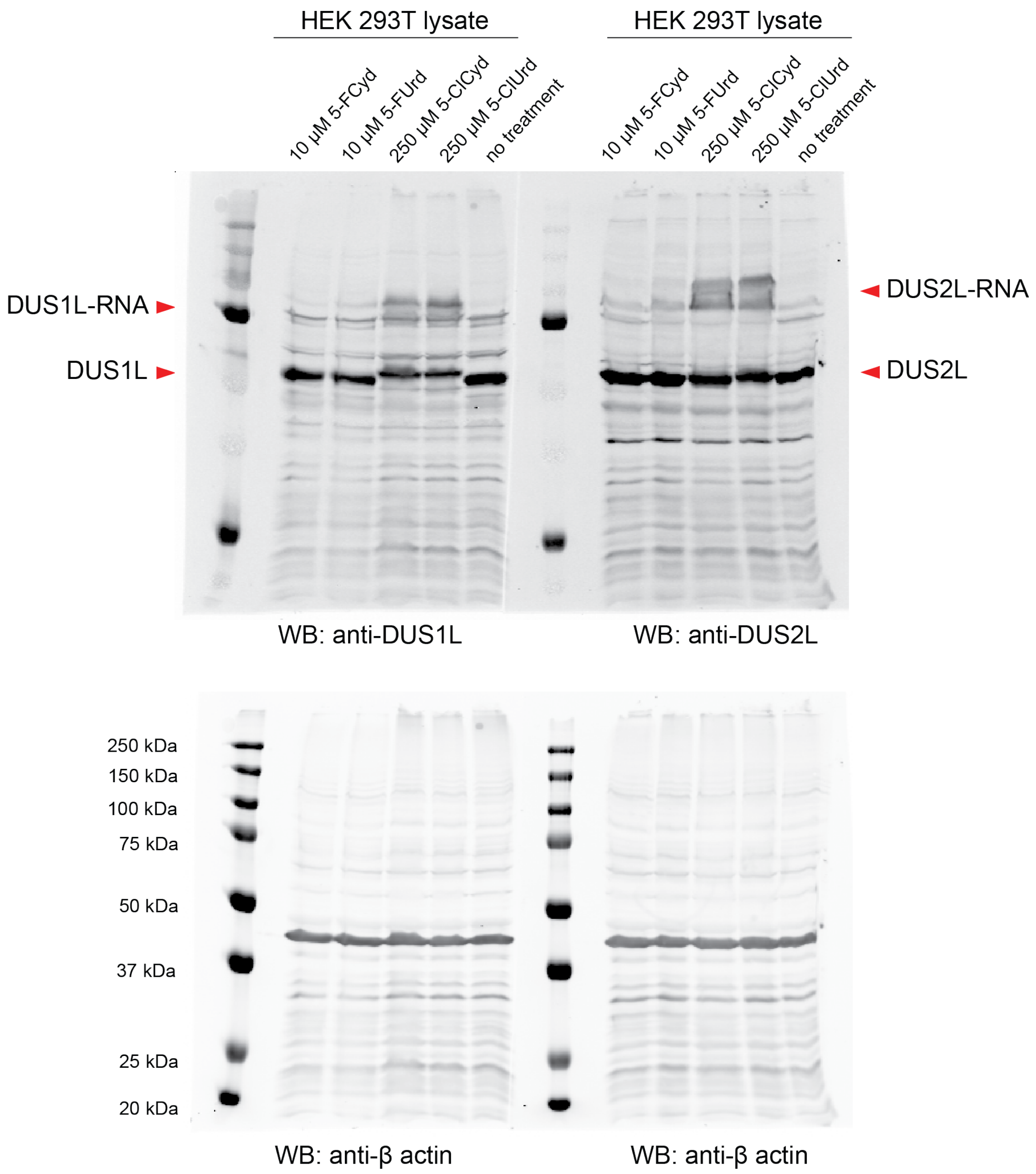


**Supplementary Figure 2.** Western blot validation of mechanism-based crosslinking between DUS1L/DUS2L and 5-halopyrimidine analogs. Full western blots for Fig. 2d and 2e. HEK293T WT cells treated with 10 µM 5-FCyd, 10 µM 5-FUrd, 250 µM 5-ClCyd, or 250 µM 5-ClUrd. Non-treated cells were used as a negative control. Both DUS1L and DUS2L were stained. β-actin was used as a loading control. Experiment was repeated 2 times independently with similar results.

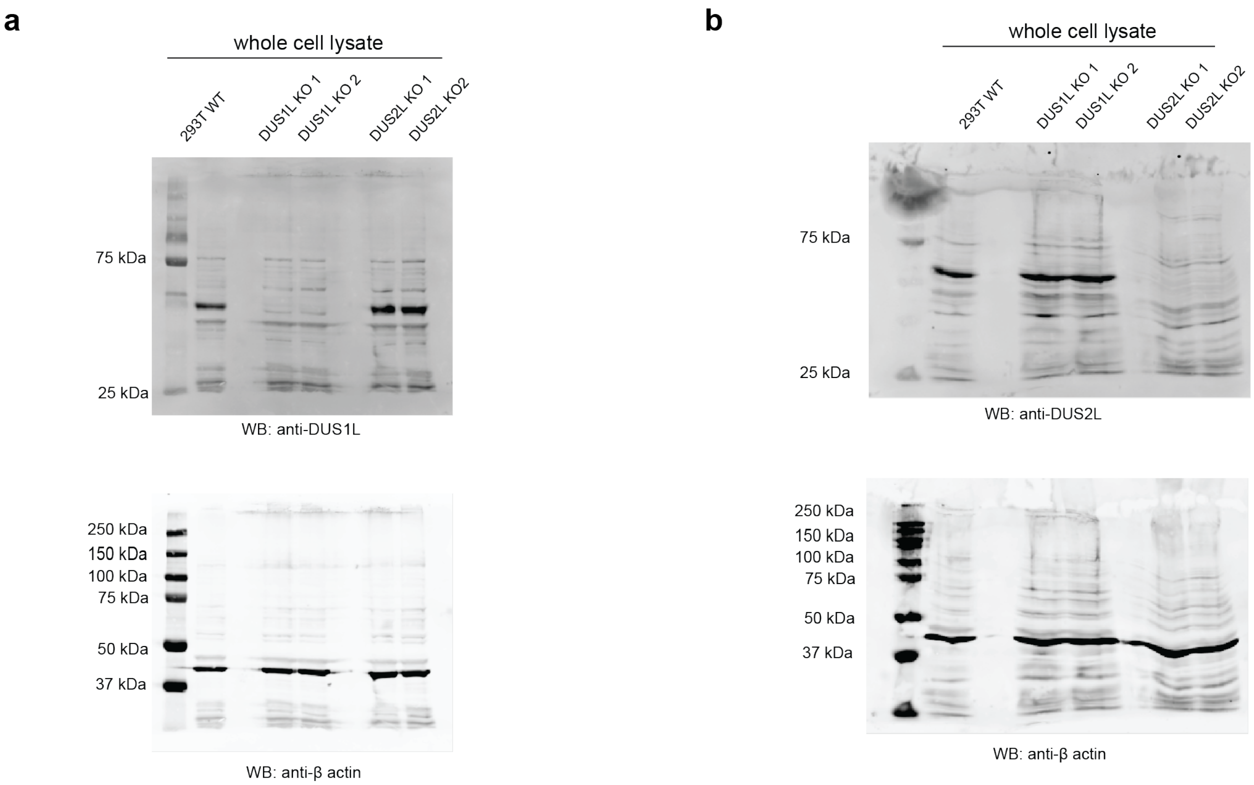


**Supplementary Figure 3.** Western blot validation of HEK293T DUS KO cell lines. **(a)** Western blot of of HEK293T WT, DUS1L KO #1, DUS1L KO #2, DUS2L KO #1, and DUS2L KO #2 using anti-DUS1L antibody. **(b)** Western blot of HEK293T WT, DUS1L KO #1, DUS1L KO #2, DUS2L KO #1, and DUS2L KO #2 using anti-DUS2L antibody. β-actin was used as a loading control. For **(a)** and **(b)** the experiments were repeated 3 times independently with similar results.

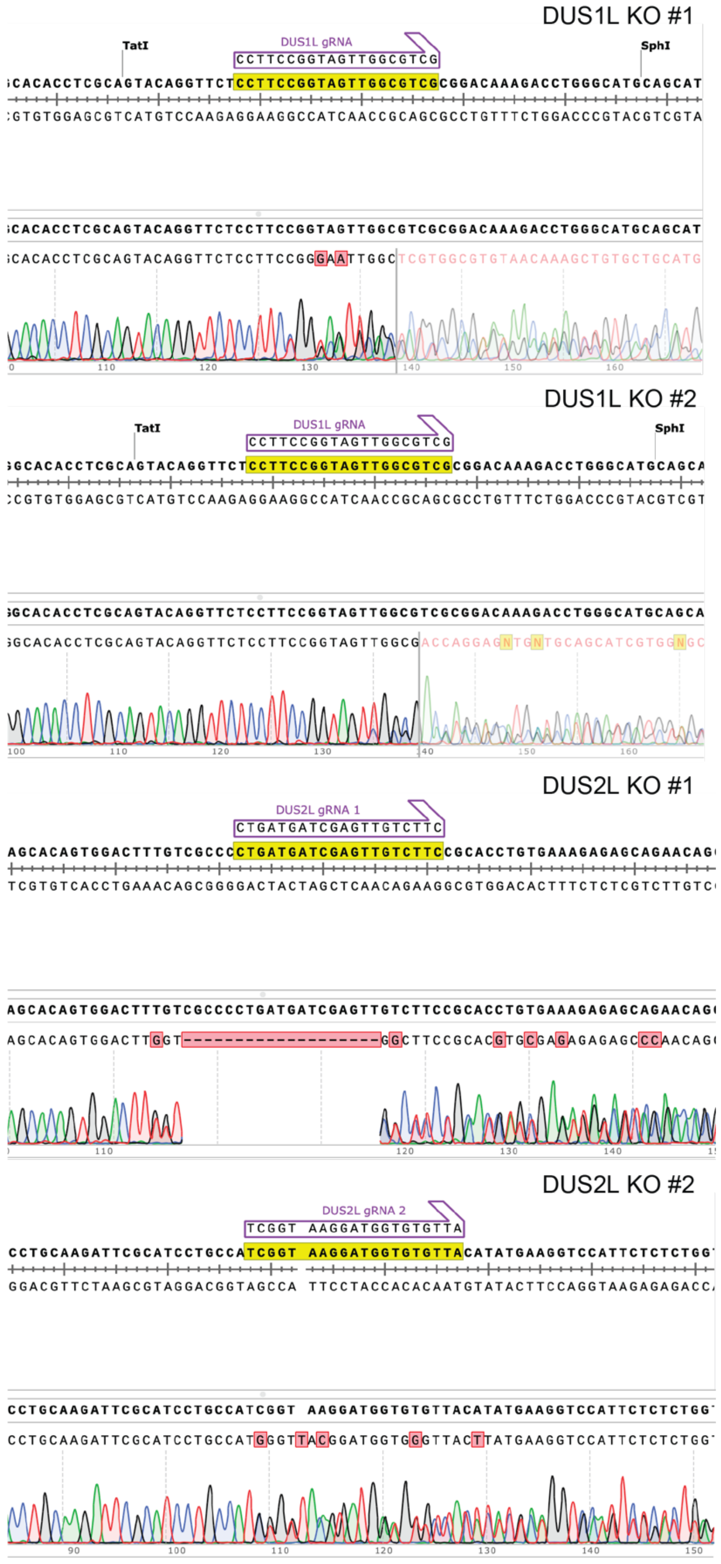


**Supplementary Figure 4.** Sanger sequencing results for genomic PCR of HEK293T DUS KO cells. The guide RNA site is highlighted.

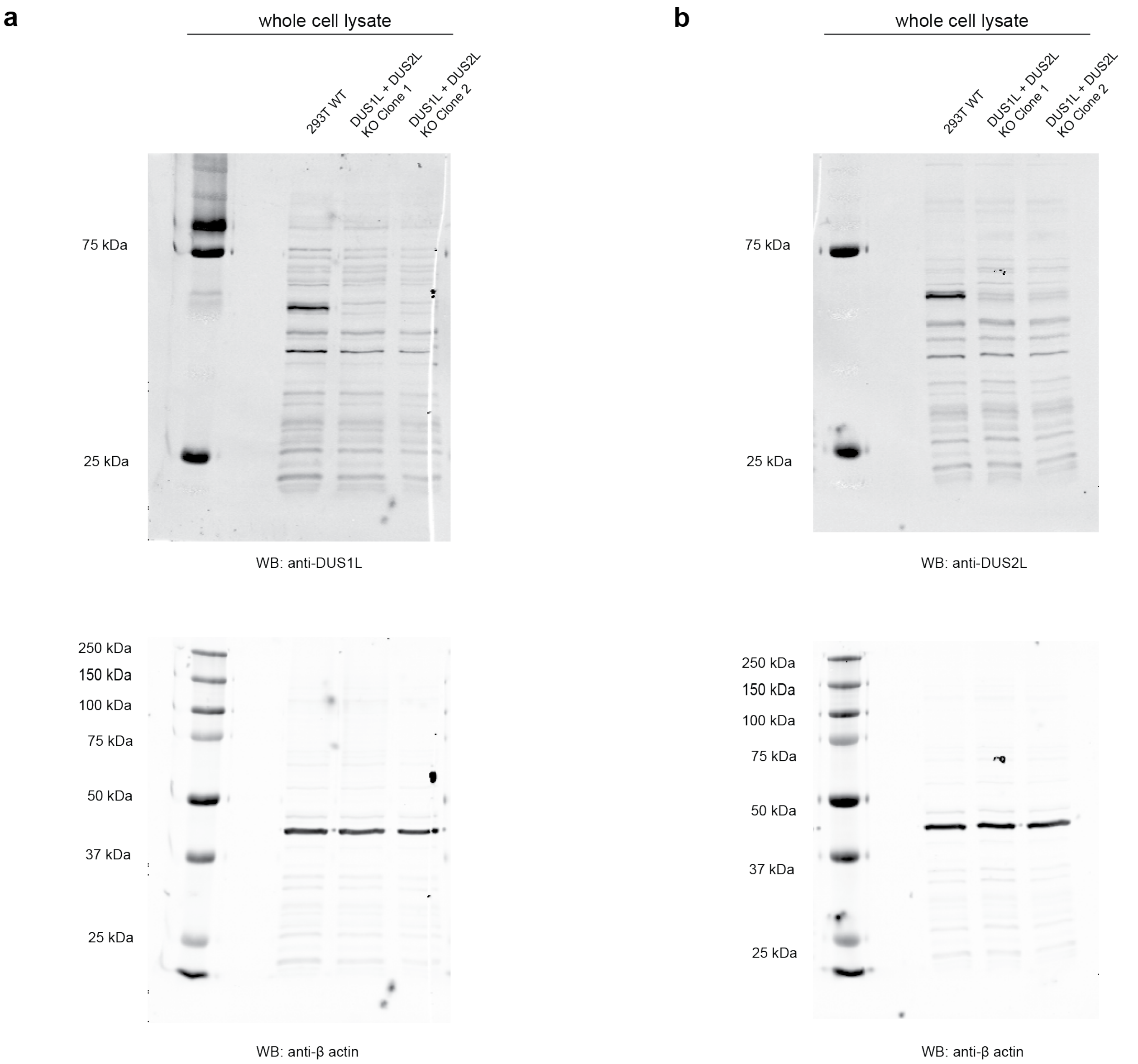


**Supplementary Figure 5.** Western blot validation of double DUS1L and DUS2L KO cell lines. **(a)** Western blot of HEK293T WT, DUS1L + 2L KO #1, and DUS1L + 2L KO #2 with anti-DUS1L antibody. **(b)** Western blot of HEK293T WT, DUS1L + 2L KO #1, and DUS1L + 2L KO #2 with anti-DUS2L antibody. β-actin was used as a loading control. For **(a)** and **(b)** the experiments were repeated 3 times independently with similar results.

DUS1L + DUS2L KO #1

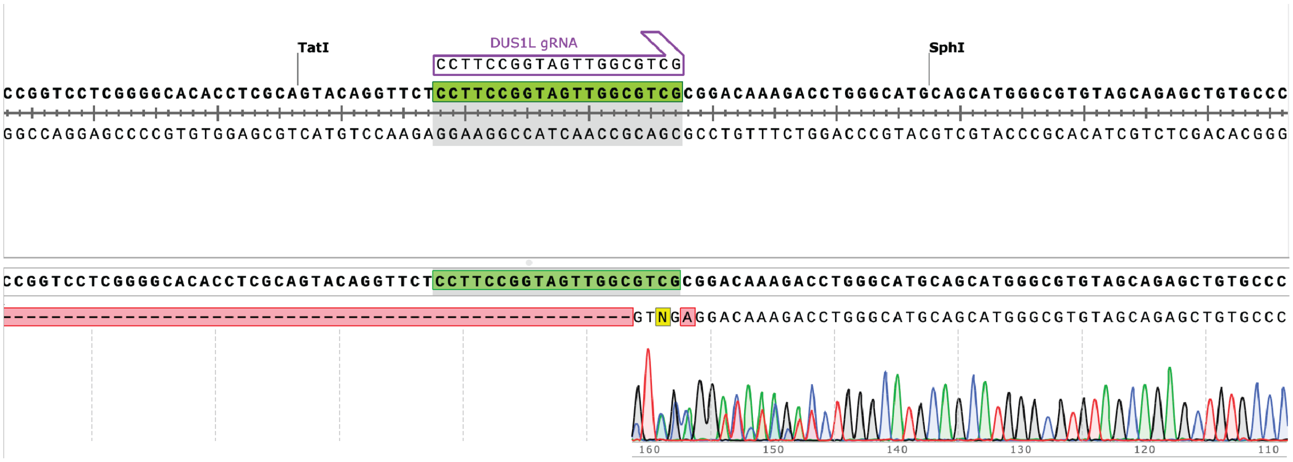


DUS1L + DUS2L KO #2

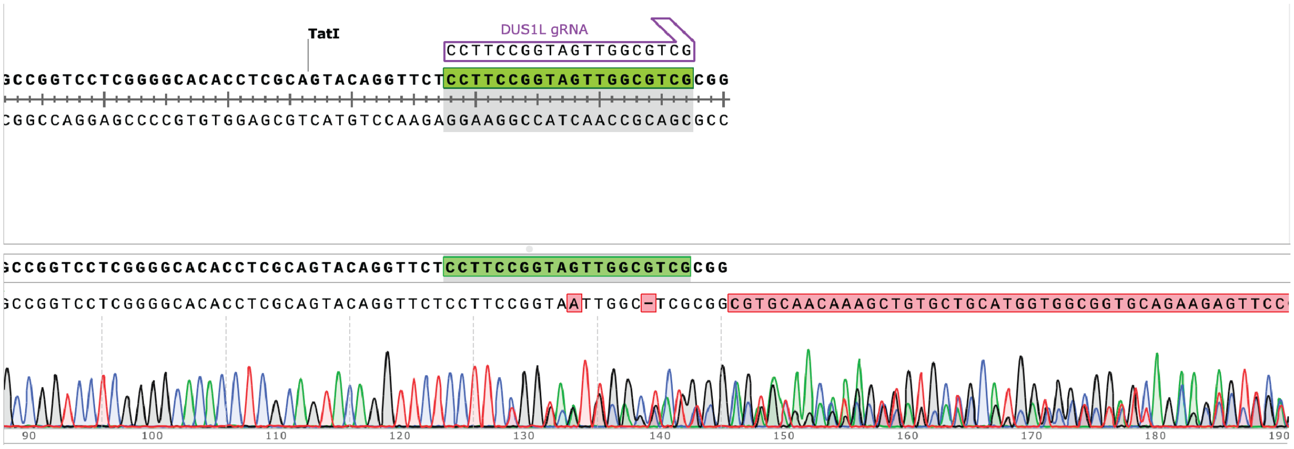


**Supplementary Figure 6.** Sanger sequencing results for genomic PCR of DUS1L + DUS2L KO HEK293T cells. Double KO line was generated by transfecting DUS1L guide RNA into DUS2L KO clone 1. The guide RNA site is highlighted.

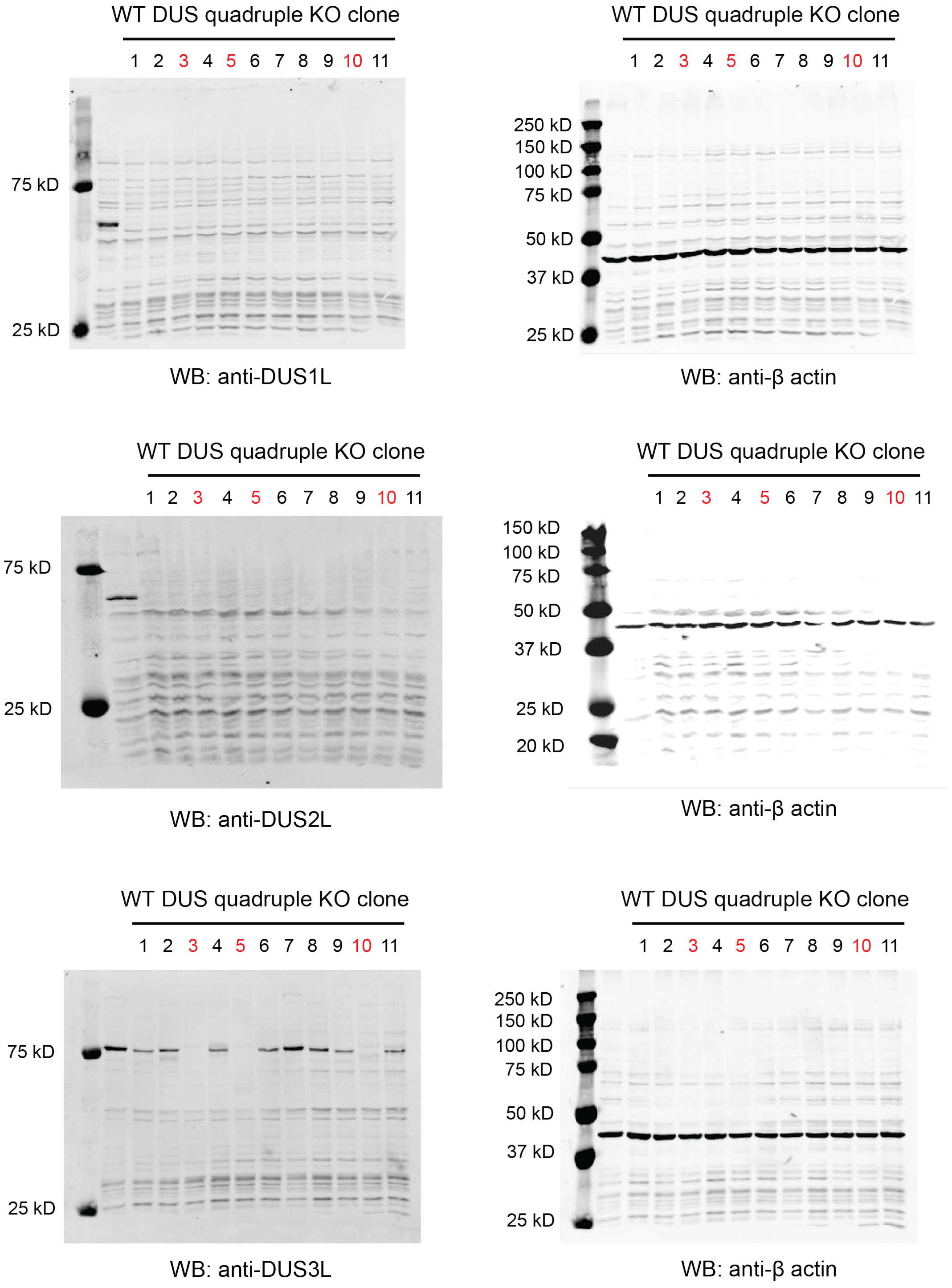


**Supplementary Figure 7.** Characterization of HEK293T DUS1L + DUS2L + DUS3L + DUS4L (ΔDUS KO) cell lines. Western blot of HEK293T ΔDUS KO. Anti-DUS1L, anti-DUS2L, and anti-DUS3L blots are shown with anti-β actin staining as a loading control. Experiments were independently repeated twice with similar results.

**a**

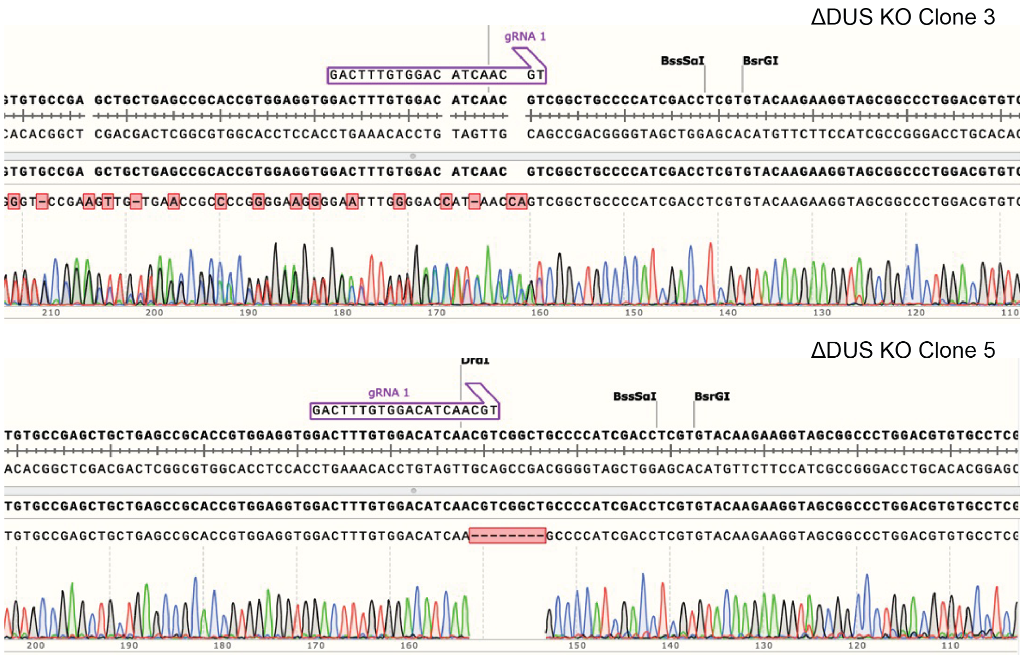


**b**

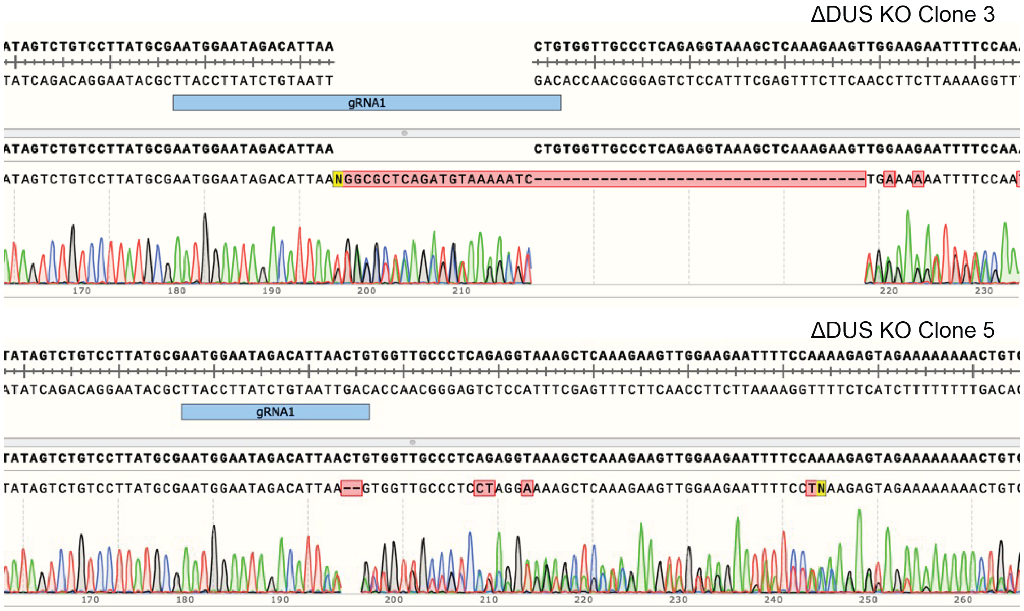


**Supplementary Figure 8.** Sanger sequencing results of genomic PCR for ΔDUS KO HEK293T cell lines. **(a)** Sanger sequencing results of genomic PCR of DUS3L in ΔDUS KO. **(b)** Sanger sequencing results of genomic PCR of DUS4L in ΔDUS KO. Guide RNA is labeled for each trace.

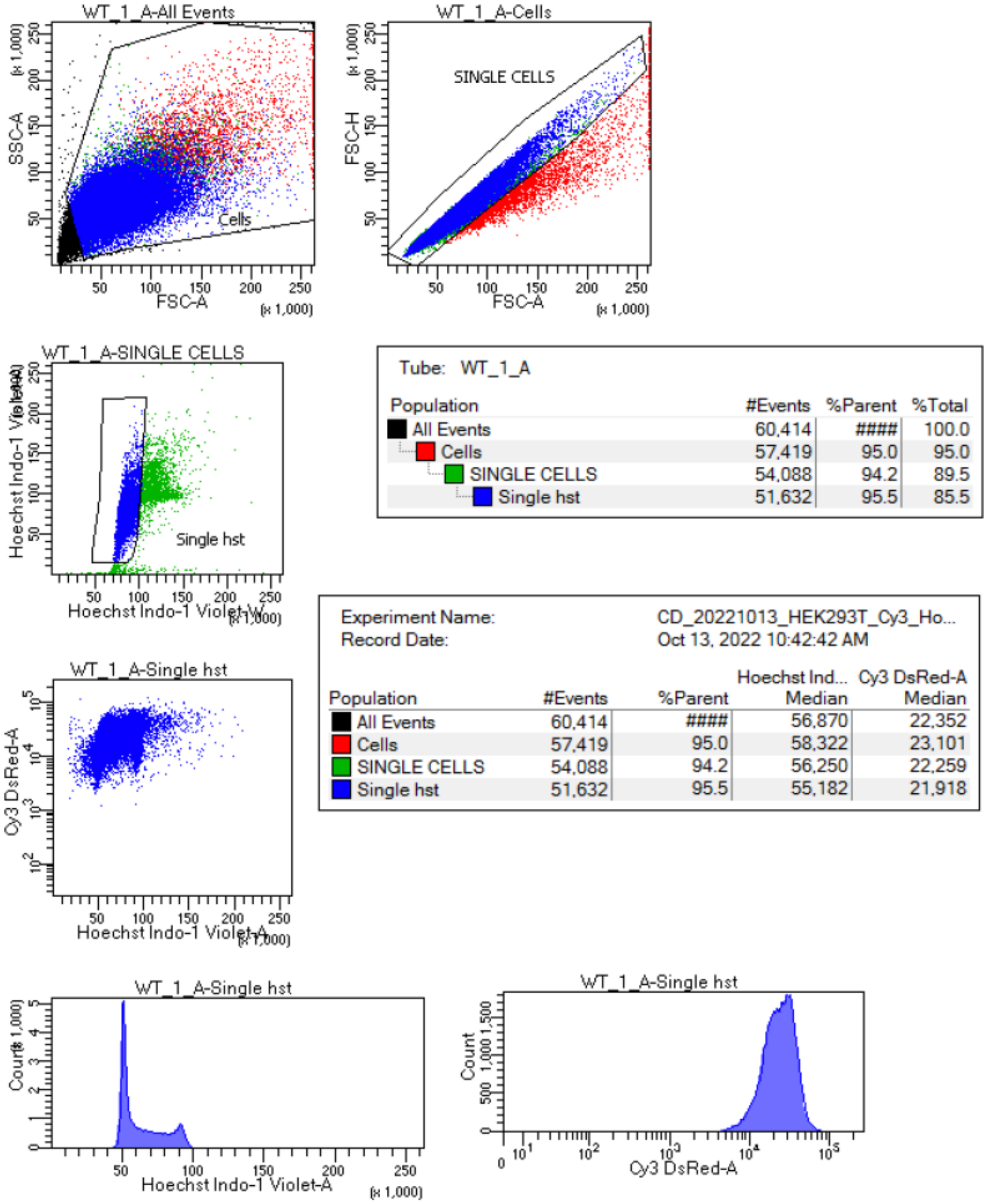


Tube: WT_1_A

| Population | #Events | %Parent | %Total |
|---|---|---|---|
| All Events | 60,414 | #### | 100.0 |
| Cells | 57,419 | 95.0 | 95.0 |
| SINGLE CELLS | 54,088 | 94.2 | 89.5 |
| Single hst | 51,632 | 95.5 | 85.5 |

Experiment Name: CD_20221013_HEK293T_Cy3_Ho...
Record Date: Oct 13, 2022 10:42:42 AM

| Population | #Events | %Parent | Hoechst Ind... Median | Cy3 DsRed-A Median |
|---|---|---|---|---|
| All Events | 60,414 | #### | 56,870 | 22,352 |
| Cells | 57,419 | 95.0 | 58,322 | 23,101 |
| SINGLE CELLS | 54,088 | 94.2 | 56,250 | 22,259 |
| Single hst | 51,632 | 95.5 | 55,182 | 21,918 |

**Supplementary Figure 9.** Example of FACS gating for OP-Puro quantitative analysis. On the SSC-A (Side Scatter-Area) vs FSC-A (Forward Scatter-Area) scatter plot, we set the primary analysis gate for only cells. On the FSC-H (Forward Scatter-Height) vs. FSC-A scatter plot, we included only single cells in the secondary analysis gate. The tertiary gate was set to exclude cellular aggregates based on a scatter plot of the Hoechst 33342-Area fluorescence intensity vs. the Hoechst 33342-Width value.

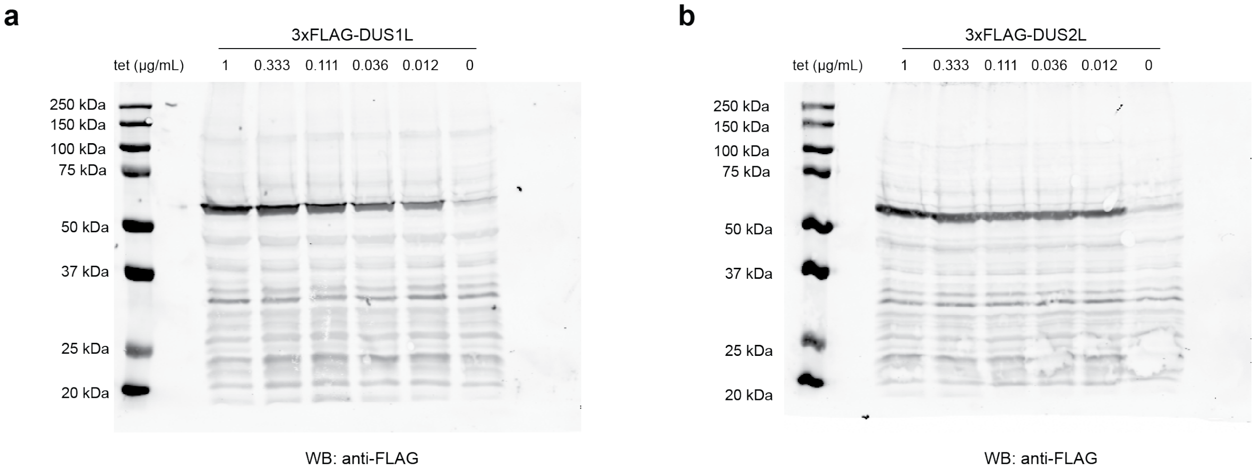


**Supplementary Figure 10.** Expression of 3xFLAG-tagged DUS1L and DUS2L in HEK293 Flp-In T-Rex stable cell lines. **(a)** Western blot expression test for Flp-In T-Rex 293 DUS1L cells. **(b)** Western blot expression test for Flp-In T-Rex 293 DUS2L cells, which were used for iCLIP. tet: tetracycline. For **(a)** and **(b)**, the experiments were repeated 3 times independently with similar results.

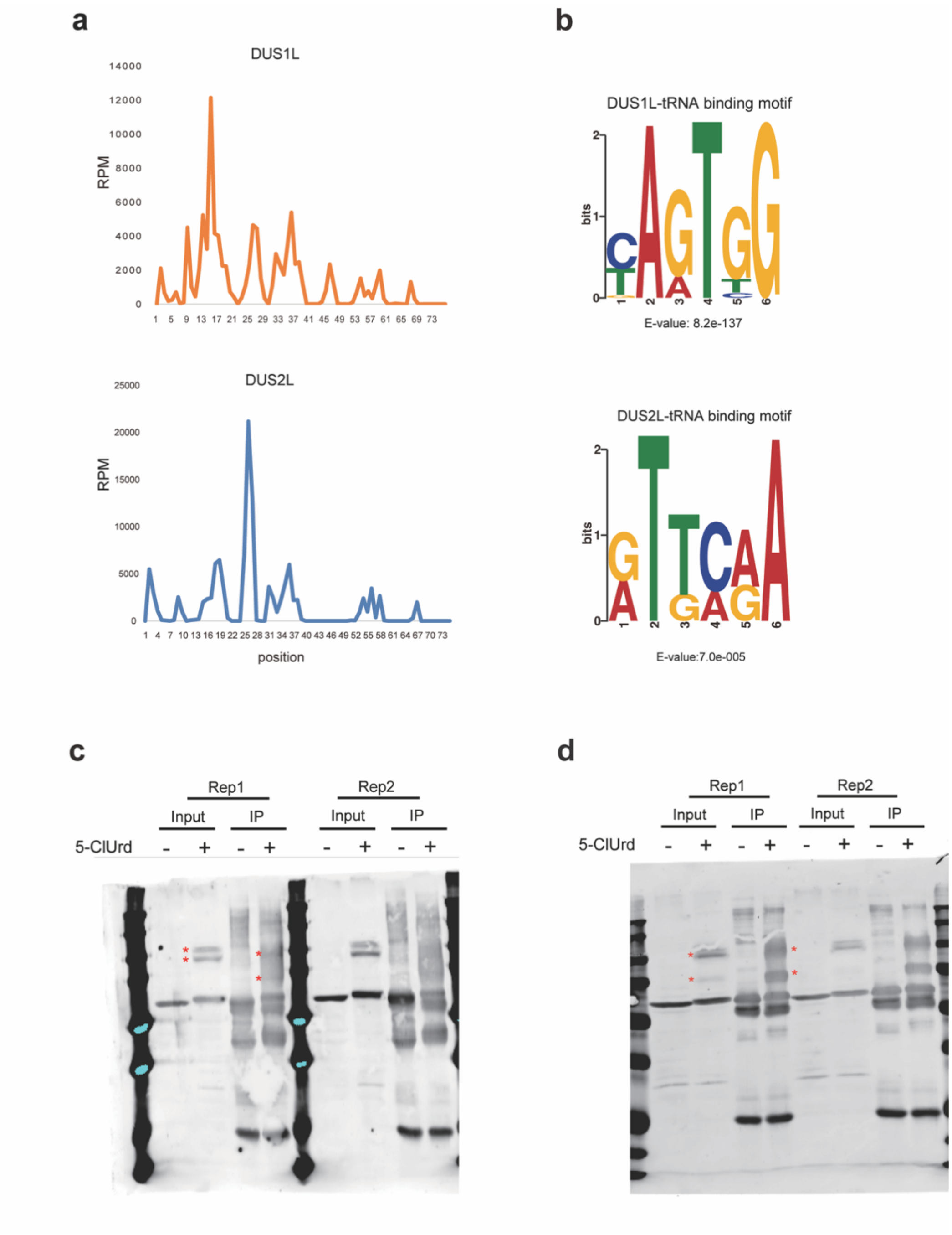


**Supplementary Figure 11.** 5-ClUrd-iCLIP data. **(a)** Coverage of all tRNA x-link peaks from DUS1L or DUS2L 5-ClUrd-iCLIP according to relative position within mature tRNA. **(b)** Consensus motif analyzed by MEME using all RNA peaks identified for DUS1L and DUS2L from 5-ClUrd-iCLIP. **(c)** Western blot of anti-FLAG 5-ClUrd-iCLIP DUS1L immunoprecipitation. **(d)** Western blot of anti-FLAG 5-ClUrd-iCLIP DUS2L immunoprecipitation. For (**c**) and (**d**), the experiments were repeated with 3 biological replicates with similar results.

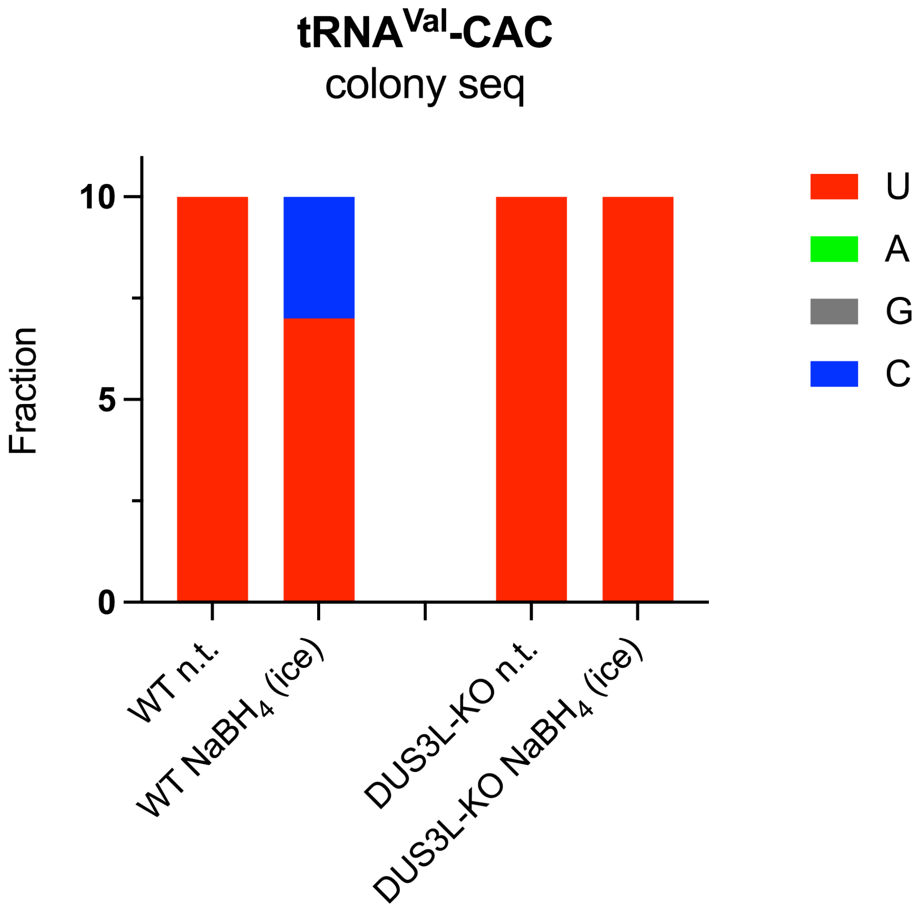


**Supplementary Figure 12.** Mutational analysis of tRNA-Val-CAC from HEK293T or DUS3L KO cells upon $NaBH_4$ reduction of RNA. Mutations at U47 in tRNA-Val-CAC were measured by colony sequencing (10 colonies per condition).

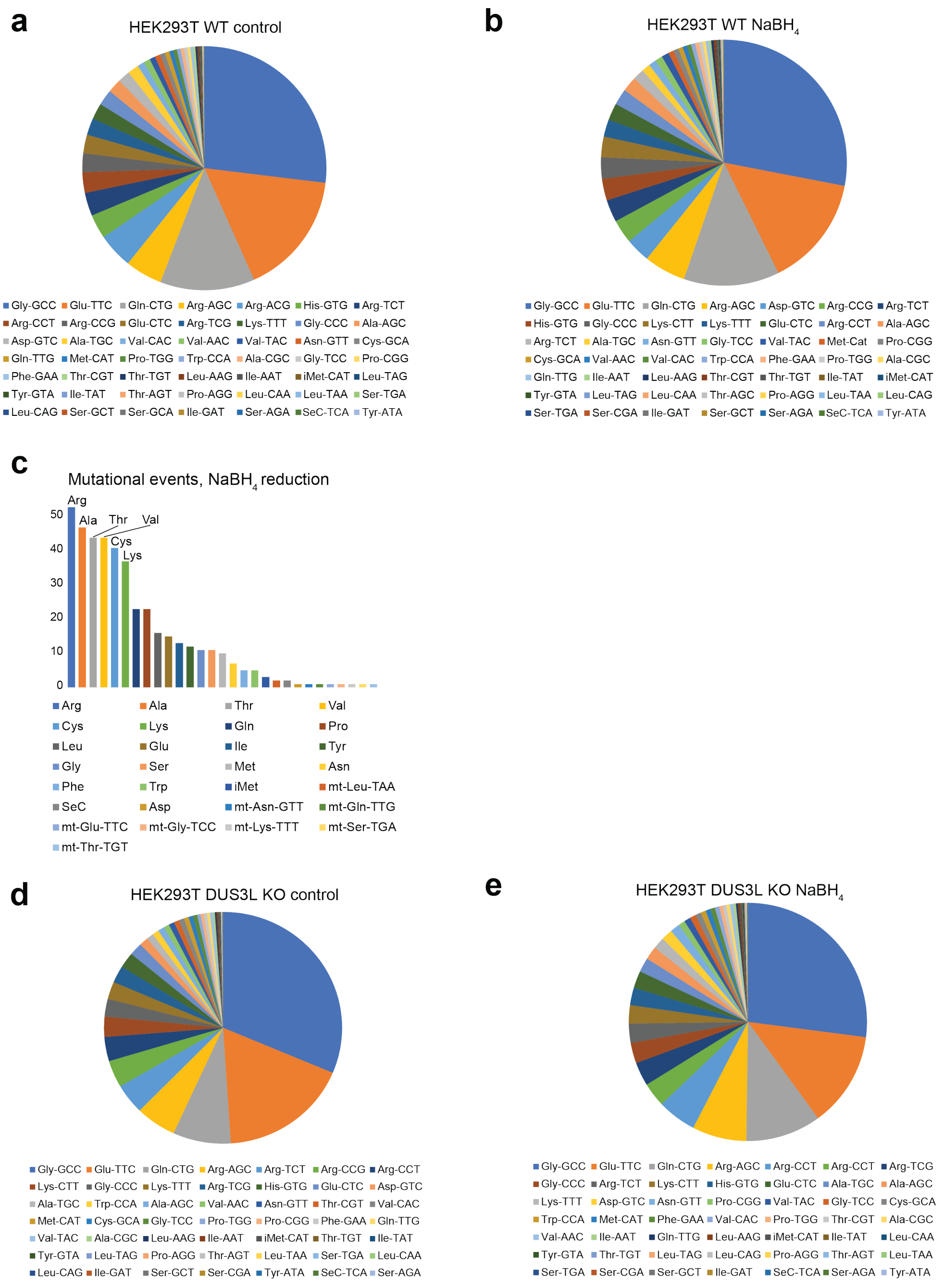


**Supplementary Figure 13.** HEK293T WT and HEK293T DUS3L KO tRNA sequencing coverage. **(a)** Fraction of mapped reads assigned to specific tRNA isodecoder families in untreated HEK293T WT cells. **(b)** Fraction of mapped reads assigned to specific tRNA isodecoder families in $NaBH_4$-treated HEK293T WT cells. Experiment was performed with three independent biological replicates for both treatment and control conditions. **(c)** Number of mutational events upon $NaBH_4$-treatment per tRNA isoacceptor family in HEK293T WT. **(d)** Fraction of mapped reads assigned to specific tRNA isodecoder families in untreated HEK293T DUS3L KO cells. **(e)** Fraction of mapped reads assigned to specific tRNA isodecoder families in $NaBH_4$-treated HEK293T DUS3L KO cells. Experiment was performed with three independent biological replicates for both treatment and control conditions.

### HEK293T DUS3L KO tRNA

**fold change ($NaBH_4$/control)**

| tRNA | 16 | 17 | 20 | 20a | 20b | 47 |
|---|---|---|---|---|---|---|
| Ala-AGC | | 17.38 | 0.84 | | | |
| Ala-CGC | | 6.29 | 0.38 | | | |
| Ala-TGC | | 3.90 | 0.49 | | | 1.54 |
| Arg-ACG | | 27.67 | 8.29 | | | |
| Arg-CCG | | | 7.89 | | | |
| Arg-CCT | | 317.74 | 5.67 | | | |
| Arg-TCG | | 69.83 | | | | |
| Arg-TCT | | 23.72 | 2.59 | | | |
| Asn-GTT | | | 8.44 | 0.94 | | |
| Asp-GTC | | 0.62 | | | | |
| Cys-GCA | | 1.48 | 1.10 | | | 1.69 |
| Gln-CTG | | 21.44 | 10.10 | | | |
| Gln-TTG | | | 3.03 | 2.10 | | |
| Glu-CTC | | | 20.07 | 4.90 | | |
| Glu-TTC | | | 6.23 | 7.22 | | |
| Gly-CCC | | | 8.00 | | | |
| Gly-GCC | | | | | | |
| Gly-TCC | | | 5.98 | | | |
| His-GTG | | | | | | |
| Ile-AAT | 0.02 | | 12.30 | 3.41 | | 2.57 |
| Ile-GAT | | | | | | |
| Ile-TAT | | | | | | 2.55 |
| Leu-AAG | | | 12.73 | | | |
| Leu-CAA | | | | | | 0.15 |
| Leu-CAG | | | | | | |
| Leu-TAA | | | | | | |
| Leu-TAG | | | 38.75 | | 2.51 | |
| Lys-CTT | 5.98 | | 5.06 | | | 0.47 |
| Lys-TTT | 9.23 | | 44.68 | | | 1.73 |
| Met-CAT | 24.08 | | | | | |
| Phe-GAA | 0.18 | 4.34 | | | | |
| Pro-AGG | | | 14.36 | | | |
| Pro-CGG | | | 11.94 | | | |
| Pro-TGG | | 3.49 | 15.05 | | | |
| SeC-TCA | | | | | | |
| Ser-AGA | | | | | | 7.78 |
| Ser-CGA | | | | | | |
| Ser-GCT | | | | | | |
| Ser-TGA | | 5.50 | | | | |
| Thr-AGT | | | | | | |
| Thr-CGT | | 11.65 | | | | 2.21 |
| Thr-TGT | | | 0.44 | 7.11 | | |
| Trp-CCA | | 9.13 | | | | |
| Tyr-ATA | | | 2.00 | | | |
| Tyr-GTA | | 1.62 | 4.21 | | | |
| Val-AAC | | 2.71 | 8.14 | | | 1.83 |
| Val-CAC | | 49.98 | 52.34 | 2.99 | | 3.13 |
| Val-TAC | | 11.78 | 23.42 | 3.74 | | |
| iMet-CAT | | | | | | 0.89 |

**$NaBH_4$ reduction**

| tRNA | 16 | 17 | 20 | 20a | 20b | 47 |
|---|---|---|---|---|---|---|
| Ala-AGC | | 2.19 | 3.92 | | | |
| Ala-CGC | | 2.26 | 4.73 | | | |
| Ala-TGC | | 2.37 | 3.95 | | | 3.04 |
| Arg-ACG | | 1.99 | 2.05 | | | |
| Arg-CCG | | 1.38 | 1.85 | | | |
| Arg-CCT | | 8.73 | 1.32 | | | |
| Arg-TCG | | 6.02 | | | | |
| Arg-TCT | | 3.86 | 1.79 | | | |
| Asn-GTT | | 1.41 | 1.03 | 6.51 | | |
| Asp-GTC | | 19.11 | | | | |
| Cys-GCA | | 1.99 | 15.78 | | | 1.78 |
| Gln-CTG | | 5.16 | 2.20 | | | |
| Gln-TTG | | | 2.25 | 2.30 | | |
| Glu-CTC | | | 1.01 | 1.29 | | |
| Glu-TTC | | | 1.97 | 1.78 | | |
| Gly-CCC | | | 1.73 | | | |
| Gly-GCC | | | | | | |
| Gly-TCC | | | 1.47 | | | |
| His-GTG | | | | | | |
| Ile-AAT | 0.40 | 1.51 | 7.78 | 3.39 | | 1.07 |
| Ile-GAT | | | | | | |
| Ile-TAT | | | | | | 1.27 |
| Leu-AAG | | | 4.91 | | 1.23 | |
| Leu-CAA | | | | | | 2.55 |
| Leu-CAG | | | 4.17 | | | |
| Leu-TAA | | | | | | |
| Leu-TAG | | | 6.23 | | 2.92 | |
| Lys-CTT | 3.57 | | 2.08 | | | 1.75 |
| Lys-TTT | 2.84 | | 1.20 | | | 4.17 |
| Met-CAT | 6.77 | | | | | |
| Phe-GAA | 2.06 | 58.95 | | | | |
| Pro-AGG | | | 3.28 | | | |
| Pro-CGG | | | 2.85 | | | |
| Pro-TGG | | 1.44 | 2.65 | | | |
| SeC-TCA | | | | | | |
| Ser-AGA | | | | | | 1.70 |
| Ser-CGA | | | | | | |
| Ser-GCT | | | | | | |
| Ser-TGA | | 21.32 | | | | |
| Thr-AGT | | | | | | |
| Thr-CGT | | 8.06 | | | | 1.21 |
| Thr-TGT | | | 9.77 | 5.65 | | |
| Trp-CCA | | 3.05 | | | | |
| Tyr-ATA | | | 15.61 | | | |
| Tyr-GTA | | 5.21 | 5.24 | | | |
| Val-AAC | | 2.04 | 3.16 | | | 2.76 |
| Val-CAC | | 1.37 | 3.28 | 2.68 | | 1.99 |
| Val-TAC | | 1.95 | 1.60 | 2.59 | | |
| iMet-CAT | | | | | | 84.38 |

**control**

| tRNA | 16 | 17 | 20 | 20a | 20b | 47 |
|---|---|---|---|---|---|---|
| Ala-AGC | | 0.13 | 4.65 | | | |
| Ala-CGC | | 0.36 | 12.33 | | | |
| Ala-TGC | | 0.61 | 8.11 | | | 1.97 |
| Arg-ACG | | 0.07 | 0.25 | | | |
| Arg-CCG | | 0.00 | 0.23 | | | |
| Arg-CCT | | 0.03 | 0.23 | | | |
| Arg-TCG | | 0.09 | | | | |
| Arg-TCT | | 0.16 | 0.69 | | | |
| Asn-GTT | | 0.00 | 0.12 | 6.95 | | |
| Asp-GTC | | 30.96 | | | | |
| Cys-GCA | | 1.35 | 14.39 | | | 1.05 |
| Gln-CTG | | 0.24 | 0.22 | | | |
| Gln-TTG | | | 0.74 | 1.09 | | |
| Glu-CTC | | | 0.05 | 0.26 | | |
| Glu-TTC | | | 0.32 | 0.25 | | |
| Gly-CCC | | | 0.22 | | | |
| Gly-GCC | | | | | | |
| Gly-TCC | | | 0.25 | | | |
| His-GTG | | | | | | |
| Ile-AAT | 19.85 | 0.00 | 0.63 | 1.00 | | 0.42 |
| Ile-GAT | | | | | | |
| Ile-TAT | | | | | | 0.50 |
| Leu-AAG | | | 0.39 | | 0.00 | |
| Leu-CAA | | | | | | 17.43 |
| Leu-CAG | | | 0.00 | | | |
| Leu-TAA | | | | | | |
| Leu-TAG | | | 0.16 | | 1.16 | |
| Lys-CTT | 0.60 | | 0.41 | | | 3.73 |
| Lys-TTT | 0.31 | | 0.03 | | | 2.40 |
| Met-CAT | 0.28 | | | | | |
| Phe-GAA | 11.59 | 13.59 | | | | |
| Pro-AGG | | | 0.23 | | | |
| Pro-CGG | | | 0.24 | | | |
| Pro-TGG | | 0.41 | 0.18 | | | |
| SeC-TCA | | | | | | |
| Ser-AGA | | | | | | 0.22 |
| Ser-CGA | | | | | | |
| Ser-GCT | | | | | | |
| Ser-TGA | | 3.88 | | | | |
| Thr-AGT | | | | | | |
| Thr-CGT | | 0.69 | | | | 0.55 |
| Thr-TGT | | | 21.97 | 0.79 | | |
| Trp-CCA | | 0.33 | | | | |
| Tyr-ATA | | | 7.82 | | | |
| Tyr-GTA | | 3.22 | 1.24 | | | |
| Val-AAC | | 0.75 | 0.39 | | | 1.50 |
| Val-CAC | | 0.03 | 0.06 | 0.90 | | 0.63 |
| Val-TAC | | 0.17 | 0.07 | 0.69 | | |
| iMet-CAT | | | | | | 95.31 |

fold change ≤ 3
fold change > 3
<1% mutation
1-5% mutation
5-10% mutation
>10% mutation

**Supplemental Figure 14.** $NaBH_4$-based mutational profiling of tRNA D sites in HEK293T DUS3L KO cell line. Analysis of mutations at canonical D positions in tRNA from DUS3L KO cells. Fold change of $NaBH_4$ versus control (untreated) is shown for each tRNA anticodon species. $NaBH_4$ and control mutational rates are shown.

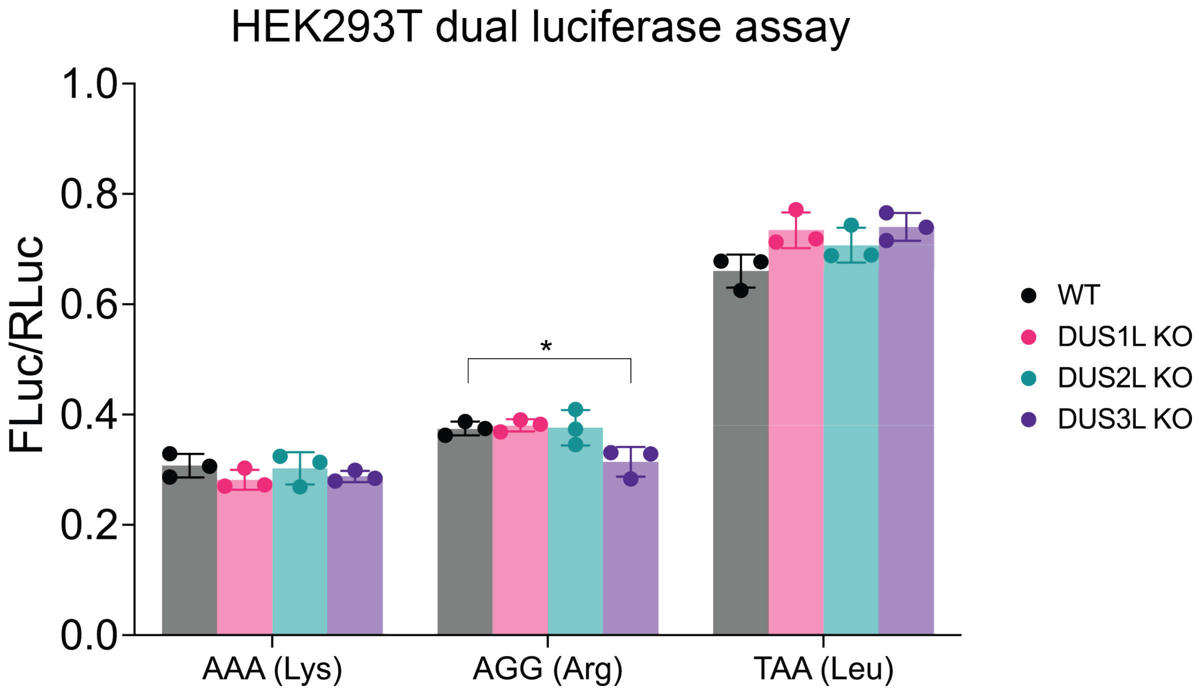


**Supplementary Figure 15.** HEK293T dual luciferase reporter data. 15 repeats of the specified codon were incorporated into the linker region between Firefly luciferase and Renilla luciferase. Three independent biological replicates were analyzed. An unpaired t-test (two-tailed) was used to measure the statistical significance *$p < 0.05$.

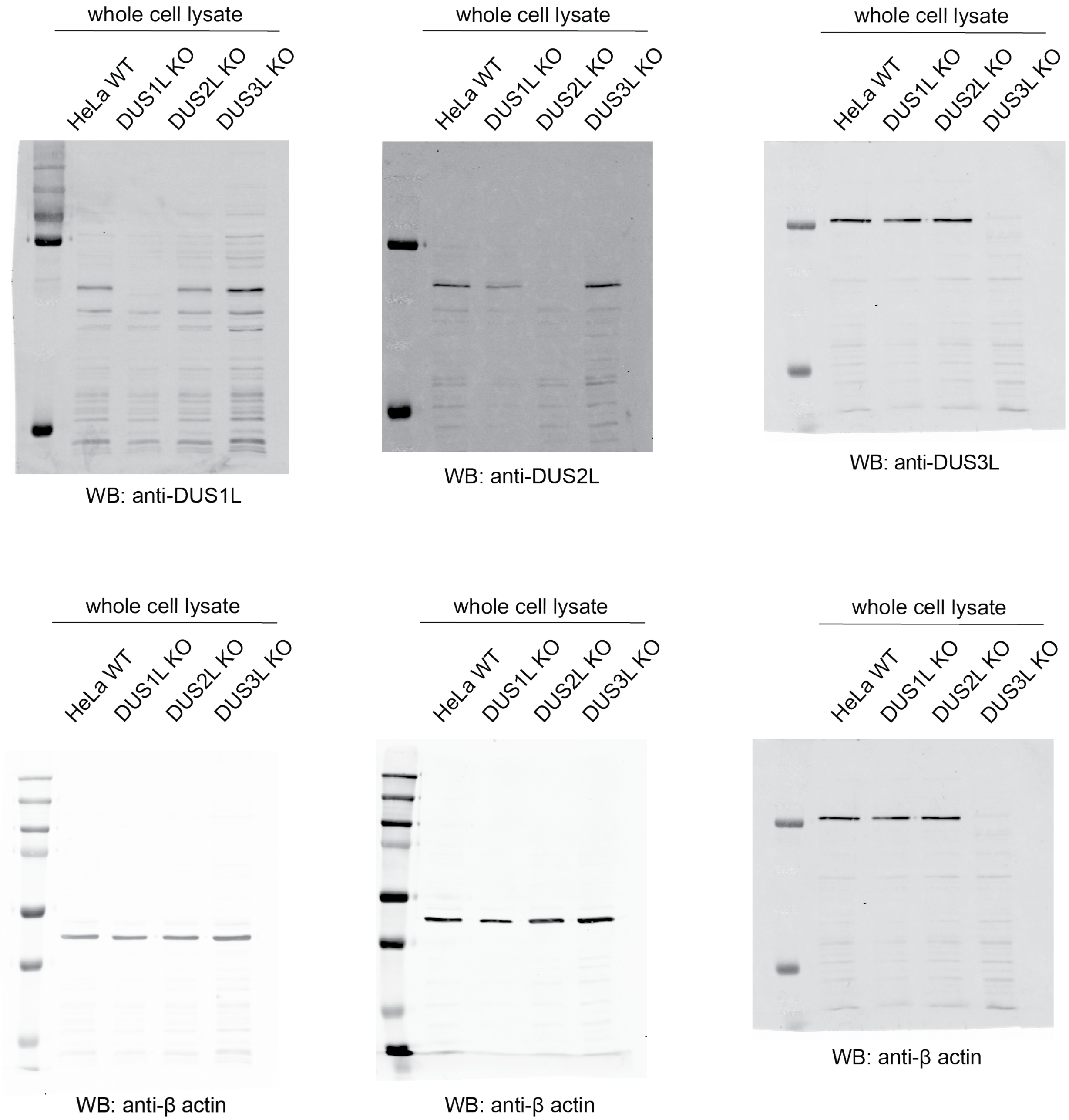


**Supplementary Figure 16.** Characterization of HeLa DUS KO cell lines. Western blot of HeLa WT, DUS1L KO, DUS2L KO, and DUS3L KO cell lines. β-actin was used as a loading control. Western blots were repeated 3 times independently with similar results.

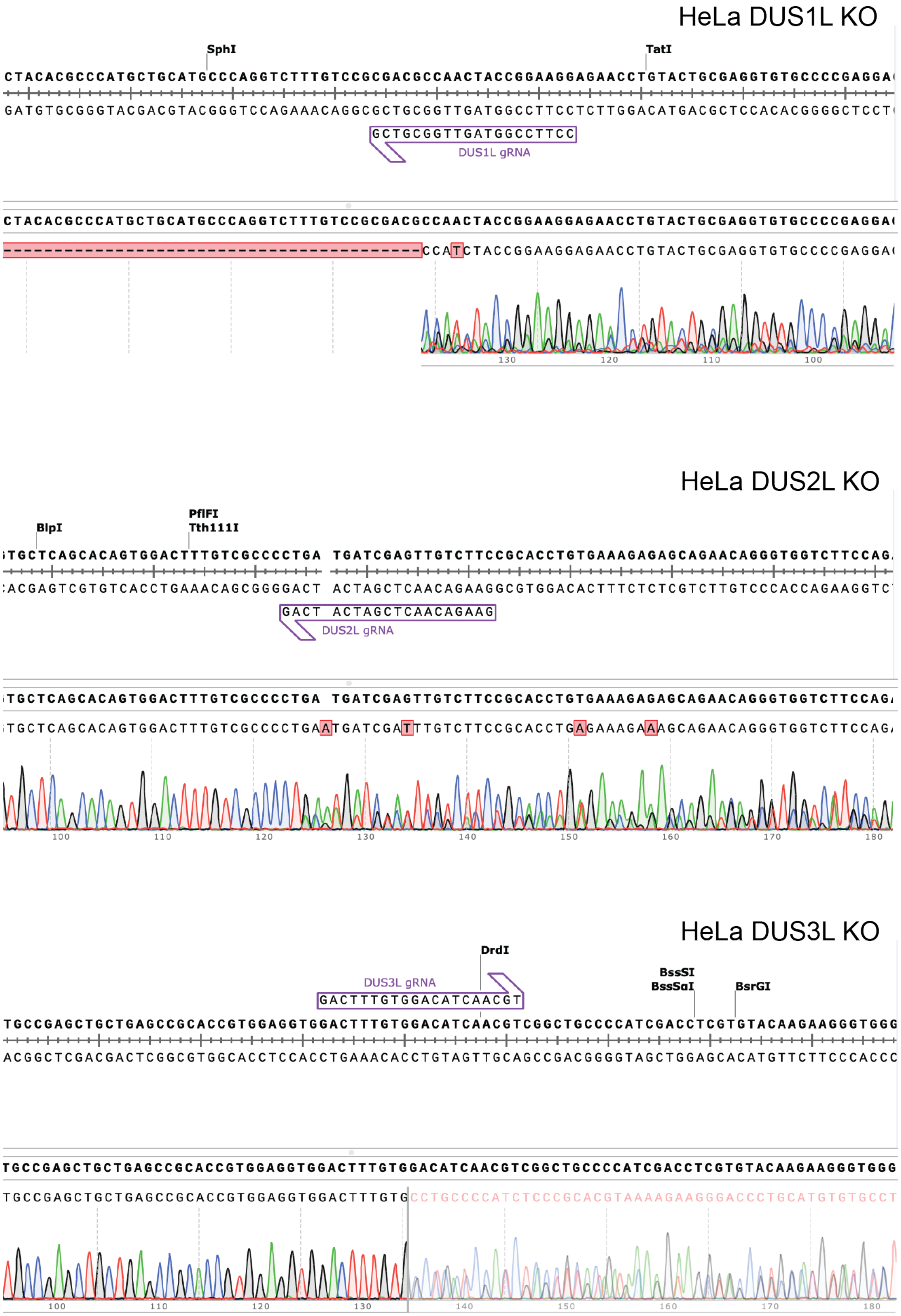


**Supplementary Figure 17.** Characterization of HeLa DUS KO cell lines. Sanger sequencing data from genomic PCR of HeLa DUS KO cell lines. Guide RNA binding sites are shown.

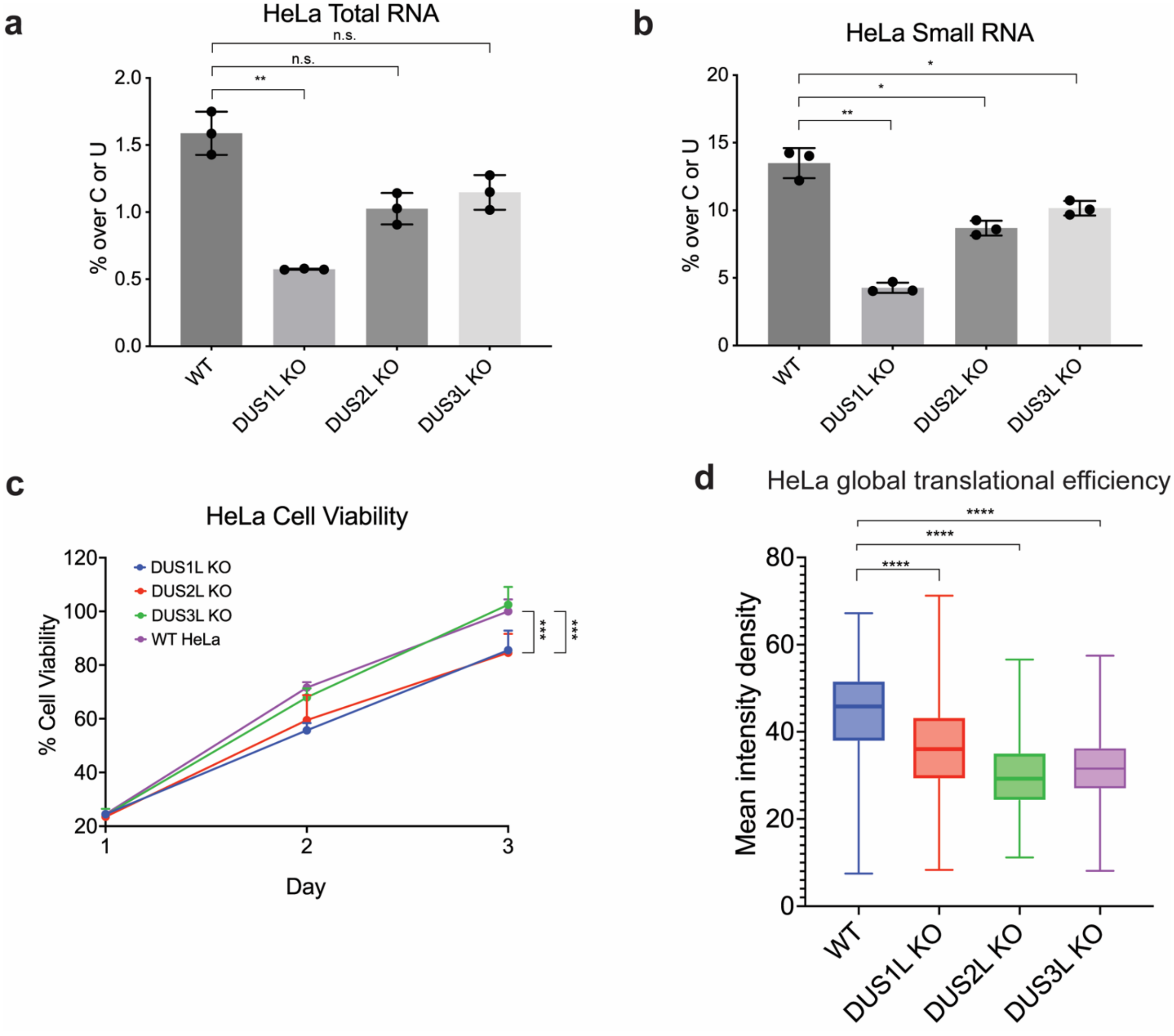


**Supplementary Figure 18.** Characterization of HeLa DUS KO cell lines. **(a)** Nucleoside LC-QQQ-MS quantitative analysis of total RNA in HeLa WT, HeLa DUS1L KO, HeLa DUS2L KO, and HeLa DUS3L KO. **(b)** Nucleoside LC-QQQ-MS quantitative analysis of small RNA in HeLa WT, HeLa DUS1L KO, HeLa DUS2L KO, and HeLa DUS3L KO. For **(a)** and **(b)** three independent biological replicates were quantified per cell line. Data is shown as mean value with s. e. m. **(c)** Cell proliferation data for HeLa WT, DUS1L KO, DUS2L KO, and DUS3L KO cell lines measured by MTS assay. Data points show mean value with s. e. m. For each day, four independent biological replicates were measured per cell line. **(d)** OP-puromycin incorporation in HeLa WT, DUS1L KO, DUS2L KO, and DUS3L KO cell lines. Incorporation was measured using fluorescence microscopy after click chemistry with Cy3-azide. Data is plotted as mean intensity density of Cy3 fluorescence from individual cells. 250 cells were quantified per cell line and was repeated independently with similar results. An unpaired t-test (two-tailed) was used to measure the statistical significance $^*p < 0.05$, $^{**}p < 0.01$, $^{***}p < 0.001$, $^{****}p < 0.0001$.

### HeLa tRNA

**fold change ($NaBH_4$/control)**

| | 16 | 17 | 20 | 20a | 20b | 47 |
|---|---|---|---|---|---|---|
| Ala-AGC | | 17.12 | 0.75 | | | 9.42 |
| Ala-CGC | | 3.16 | 0.71 | | | 10.33 |
| Ala-TGC | | 2.77 | 0.80 | | | 1.63 |
| Arg-ACG | | 37.21 | 3.56 | | | 2.21 |
| Arg-CCG | | | 20.38 | | | |
| Arg-CCT | | 138.16 | 12.78 | | | |
| Arg-TCG | | | | | | |
| Arg-TCT | | 53.14 | 3.38 | | | 5.89 |
| Asn-GTT | | | | 1.56 | | |
| Asp-GTC | | | | | | |
| Cys-GCA | | 2.52 | 1.64 | | | 15.05 |
| Gln-CTG | | 6.81 | 5.23 | 7.30 | | 0.10 |
| Gln-TTG | | | 4.81 | 1.43 | | |
| Glu-CTC | | | | | | |
| Glu-TTC | | 1.31 | 5.14 | 5.73 | | |
| Gly-CCC | | | 5.01 | | | |
| Gly-GCC | | 8.16 | | | | |
| Gly-TCC | | | | | | |
| His-GTG | | | | | | |
| Ile-AAT | 0.67 | | 2.05 | 1.97 | | 3.76 |
| Ile-GAT | | | | | | |
| Ile-TAT | | | 1.85 | | | |
| Leu-AAG | | | 18.26 | | | |
| Leu-CAA | | | | | | |
| Leu-CAG | | | | | | |
| Leu-TAA | | | | | | |
| Leu-TAG | | | 16.61 | | | |
| Lys-CTT | 6.12 | | 3.75 | | | 5.46 |
| Lys-TTT | 7.34 | 2.32 | 8.38 | | | 2.81 |
| Met-CAT | 14.25 | | | | | 6.24 |
| Phe-GAA | 1.36 | | | | | 8.50 |
| Pro-AGG | | | 11.68 | | | 6.45 |
| Pro-CGG | | | 10.12 | | | 8.40 |
| Pro-TGG | | 1.87 | 6.83 | | | 6.34 |
| SeC-TCA | | | | | | |
| Ser-AGA | | | | | | |
| Ser-CGA | | | | | | |
| Ser-GCT | | | | >4.58 | | |
| Ser-TGA | | | | | | |
| Thr-AGT | | | | | | 5.73 |
| Thr-CGT | | 5.66 | | 1.05 | | 12.42 |
| Thr-TGT | | | 0.53 | | | 6.23 |
| Trp-CCA | | | | | | |
| Tyr-ATA | | | 1.50 | | | |
| Tyr-GTA | | | 1.17 | | | 5.07 |
| Val-AAC | | 2.09 | 2.73 | | | 16.86 |
| Val-CAC | | 7.27 | 4.24 | 2.16 | | 19.00 |
| Val-TAC | | | | 2.91 | | 10.32 |
| iMet-CAT | | | | | | 3.57 |

**$NaBH_4$ reduction**

| | 16 | 17 | 20 | 20a | 20b | 47 |
|---|---|---|---|---|---|---|
| Ala-AGC | | 1.78 | 2.18 | | | 3.49 |
| Ala-CGC | | 1.41 | 7.86 | | | 6.99 |
| Ala-TGC | | 1.73 | 5.39 | | | 4.46 |
| Arg-ACG | | 1.68 | 1.06 | | | 1.18 |
| Arg-CCG | | 1.62 | 1.40 | | | |
| Arg-CCT | | 2.61 | 1.49 | | | |
| Arg-TCG | | 2.39 | | | | |
| Arg-TCT | | 2.34 | 1.32 | | | 3.20 |
| Asn-GTT | | 1.34 | | 13.72 | | |
| Asp-GTC | | | | | | |
| Cys-GCA | | 1.62 | 14.62 | | | 15.52 |
| Gln-CTG | | 2.01 | 1.70 | 1.30 | | 1.85 |
| Gln-TTG | | | 1.08 | 1.82 | | |
| Glu-CTC | | | | | | |
| Glu-TTC | | 3.89 | 1.02 | 1.38 | | |
| Gly-CCC | | | 1.43 | | | |
| Gly-GCC | | 1.36 | | | | |
| Gly-TCC | | | | | | |
| His-GTG | | | | | | |
| Ile-AAT | 16.17 | | 26.39 | 1.68 | | 1.47 |
| Ile-GAT | | | | | | |
| Ile-TAT | | | 26.19 | | | |
| Leu-AAG | | | 3.86 | | | |
| Leu-CAA | | | | | | |
| Leu-CAG | | | | | | |
| Leu-TAA | | | | | | |
| Leu-TAG | | | 3.02 | | | |
| Lys-CTT | 2.58 | | 1.45 | | | 2.78 |
| Lys-TTT | 2.46 | 1.23 | 1.46 | | | 2.79 |
| Met-CAT | 8.21 | | | | | 3.33 |
| Phe-GAA | 7.00 | | | | | 7.09 |
| Pro-AGG | | | 2.23 | | | 5.35 |
| Pro-CGG | | | 2.08 | | | 6.05 |
| Pro-TGG | | 1.05 | 1.63 | | | 4.43 |
| SeC-TCA | | | | | | |
| Ser-AGA | | | | | | |
| Ser-CGA | | | | | | |
| Ser-GCT | | | | 7.32 | | |
| Ser-TGA | | | | | | |
| Thr-AGT | | | | | | 7.96 |
| Thr-CGT | | 5.03 | | 91.21 | | 5.11 |
| Thr-TGT | | | 8.90 | | | 2.70 |
| Trp-CCA | | | | | | |
| Tyr-ATA | | | 42.86 | | | |
| Tyr-GTA | | | 67.43 | | | 1.97 |
| Val-AAC | | 1.69 | 1.72 | | | 26.67 |
| Val-CAC | | 0.86 | 1.12 | 1.88 | | 25.56 |
| Val-TAC | | | 2.04 | 1.72 | | 6.33 |
| iMet-CAT | | | | | | 1.38 |

**control**

| | 16 | 17 | 20 | 20a | 20b | 47 |
|---|---|---|---|---|---|---|
| Ala-AGC | | 0.10 | 2.92 | | | 0.37 |
| Ala-CGC | | 0.44 | 11.10 | | | 0.68 |
| Ala-TGC | | 0.63 | 6.76 | | | 2.75 |
| Arg-ACG | | 0.05 | 0.30 | | | 0.54 |
| Arg-CCG | | 0.00 | 0.07 | | | |
| Arg-CCT | | 0.02 | 0.12 | | | |
| Arg-TCG | | 0.00 | | | | |
| Arg-TCT | | 0.04 | 0.39 | | | 0.54 |
| Asn-GTT | | 0.00 | | 8.79 | | |
| Asp-GTC | | | | | | |
| Cys-GCA | | 0.64 | 8.92 | | | 1.03 |
| Gln-CTG | | 0.29 | 0.33 | 0.18 | | 18.03 |
| Gln-TTG | | | 0.22 | 1.27 | | |
| Glu-CTC | | | | | | |
| Glu-TTC | | 2.97 | 0.20 | 0.24 | | |
| Gly-CCC | | | 0.29 | | | |
| Gly-GCC | | 0.17 | | | | |
| Gly-TCC | | | | | | |
| His-GTG | | | | | | |
| Ile-AAT | 24.09 | | 12.90 | 0.85 | | 0.39 |
| Ile-GAT | | | | | | |
| Ile-TAT | | | 14.14 | | | |
| Leu-AAG | | | 0.21 | | | |
| Leu-CAA | | | | | | |
| Leu-CAG | | | | | | |
| Leu-TAA | | | | | | |
| Leu-TAG | | | 0.18 | | | |
| Lys-CTT | 0.42 | | 0.39 | | | 0.51 |
| Lys-TTT | 0.34 | 0.53 | 0.17 | | | 0.99 |
| Met-CAT | 0.58 | | | | | 0.53 |
| Phe-GAA | 5.17 | | | | | 0.83 |
| Pro-AGG | | | 0.19 | | | 0.83 |
| Pro-CGG | | | 0.21 | | | 0.72 |
| Pro-TGG | | 0.56 | 0.24 | | | 0.70 |
| SeC-TCA | | | | | | |
| Ser-AGA | | | | | | |
| Ser-CGA | | | | | | |
| Ser-GCT | | | | 0.00 | | |
| Ser-TGA | | | | | | |
| Thr-AGT | | | | | | 1.39 |
| Thr-CGT | | 0.89 | | 86.49 | | 0.41 |
| Thr-TGT | | | 16.92 | | | 0.43 |
| Trp-CCA | | | | | | |
| Tyr-ATA | | | 28.57 | | | |
| Tyr-GTA | | | 57.66 | | | 0.39 |
| Val-AAC | | 0.81 | 0.63 | | | 1.58 |
| Val-CAC | | 0.12 | 0.26 | 0.87 | | 1.35 |
| Val-TAC | | | 0.00 | 0.59 | | 0.61 |
| iMet-CAT | | | | | | 0.39 |

fold change ≤ 3
fold change > 3

<1% mutation
1-5% mutation
5-10% mutation
>10% mutation

**Supplementary Figure 19.** $NaBH_4$-based mutational profiling of tRNA D sites in HeLa cells. **(a)** Analysis of mutations at canonical D positions in tRNA from HeLa cells. Fold change of $NaBH_4$ versus control (untreated) is shown for each tRNA anticodon species. $NaBH_4$ and control mutational rates are shown.

**a**

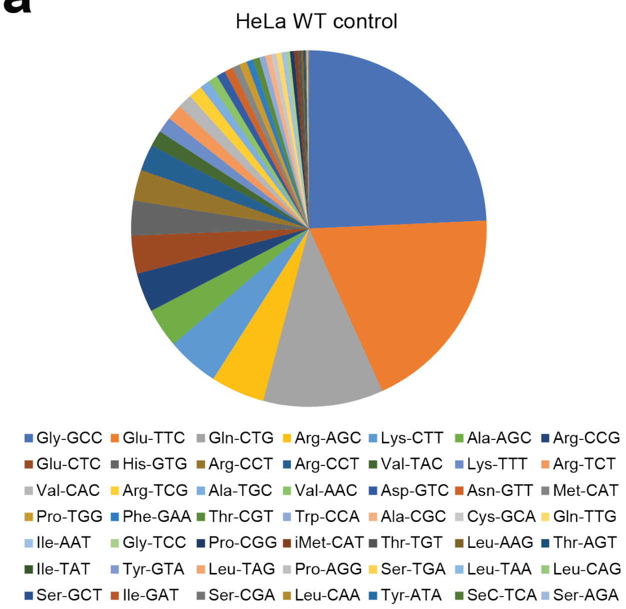


**b**

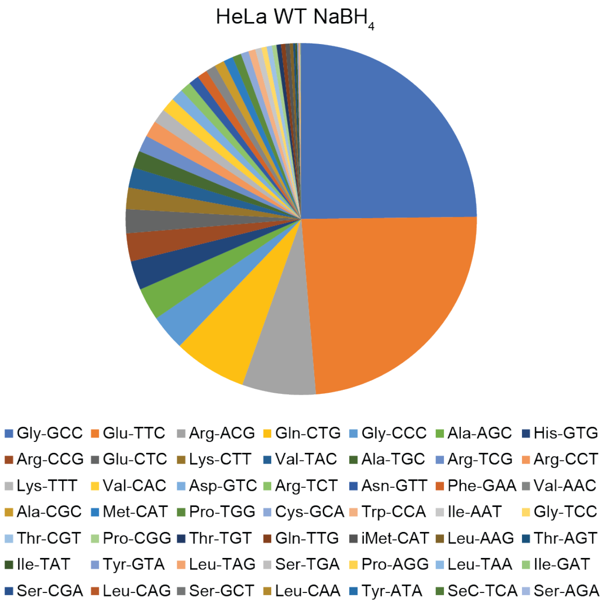


**Supplementary Figure 20.** HeLa tRNA sequencing coverage. **(a)** Fraction of mapped reads assigned to specific tRNA isodecoder families in untreated HeLa WT cells. **(b)** Fraction of mapped reads assigned to specific tRNA isodecoder families in $NaBH_4$-treated HeLa WT cells. Experiment was performed with three independent biological replicates for both treatment and control conditions.

a

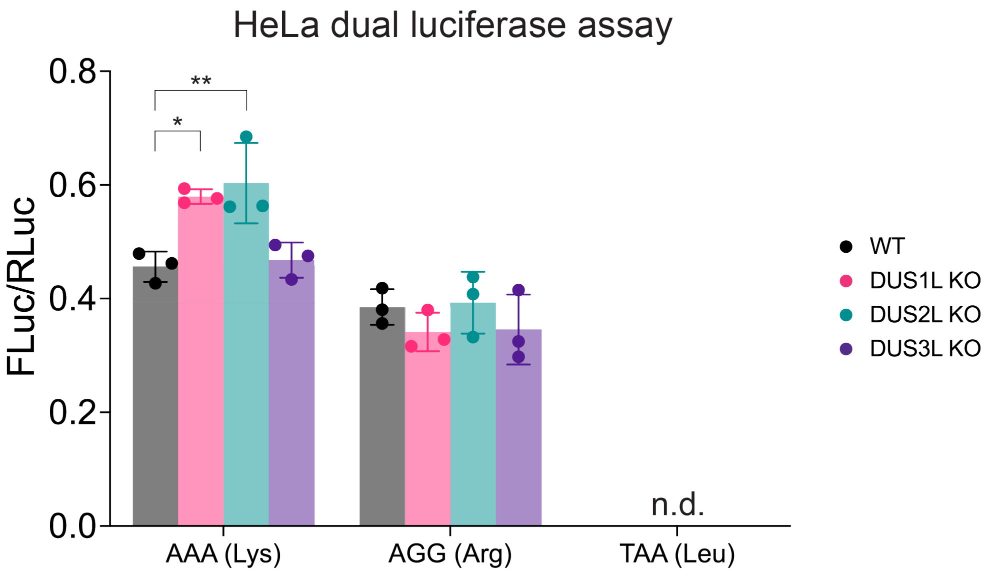


b

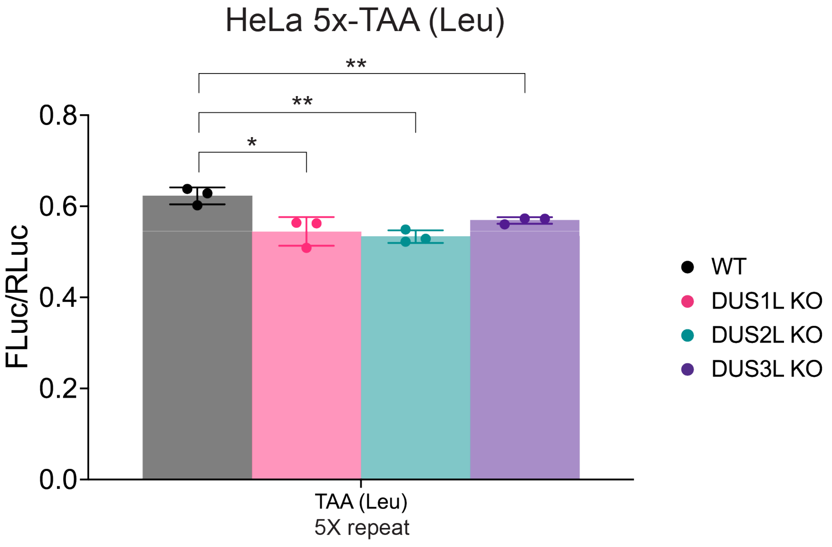


**Supplementary Figure 21.** Dual luciferase reporter data from HeLa cells. **(a)** 15 repeats of the specified codon were incorporated into the linker region between Firefly luciferase and *Renilla* luciferase; n.d. non-detectable. **(b)** For TAA (Leu), 5 repeats were used. Three independent biological replicates were analyzed. An unpaired t-test (two-tailed) was used to measure the statistical significance *p < 0.05 and **p < 0.01.

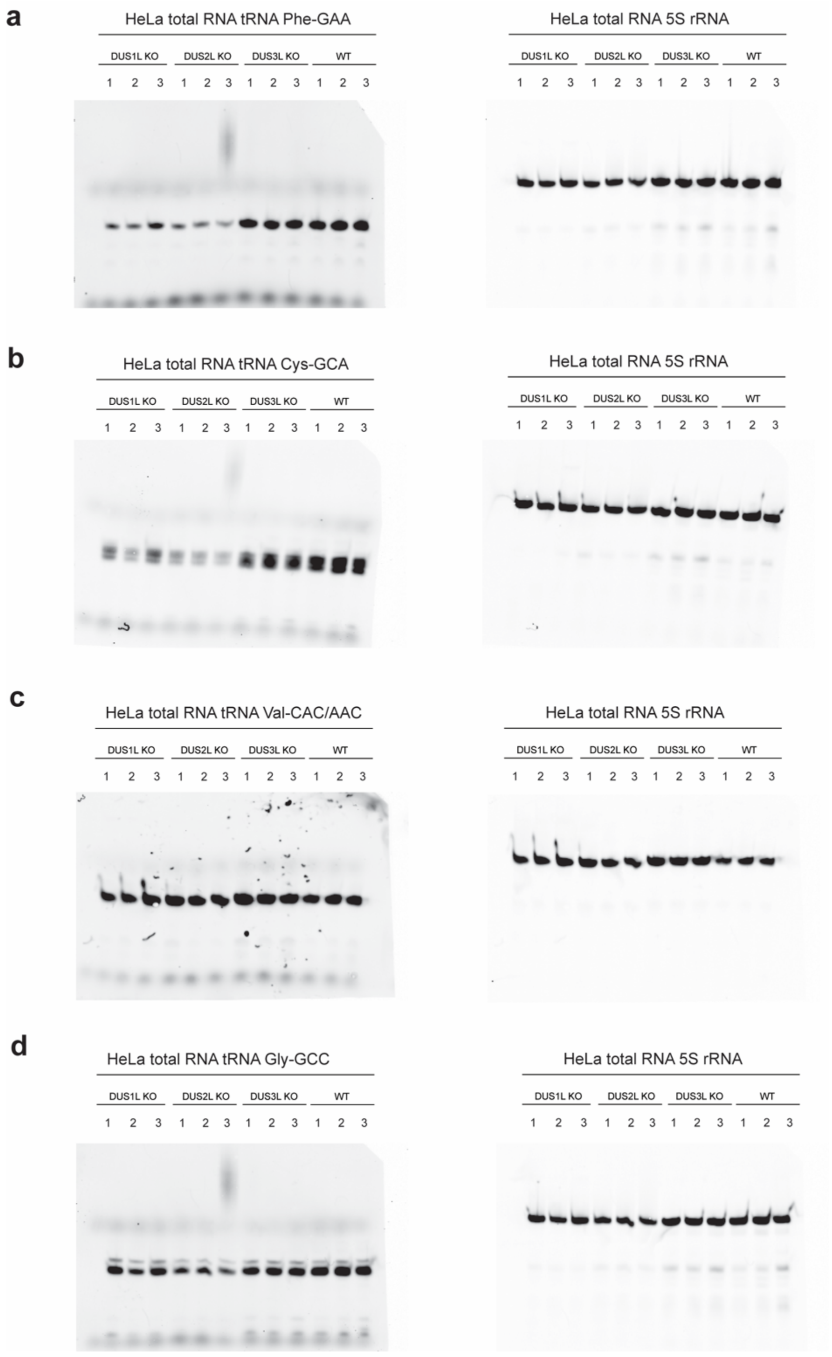


**Supplementary Figure 22.** Northern blot analysis of tRNA abundance in HeLa cells. **(a)** Total RNA from DUS1L KO, DUS2L KO, DUS3L KO, and WT HeLa cells was probed for tRNA-Phe-GAA using 5S rRNA as a loading control. **(b)** Total RNA from DUS1L KO, DUS2L KO, DUS3L KO, and WT HeLa cells was probed for tRNA-Cys-GCA using 5S rRNA as a loading control. **(c)** Total RNA from DUS1L KO, DUS2L KO, DUS3L KO, and WT HeLa cells was probed for tRNA-Val-mAC using 5S rRNA as a loading control. **(d)** Total RNA from DUS1L KO, DUS2L KO, DUS3L KO, and WT HeLa cells was probed for tRNA-Gly-GCC using 5S rRNA as a loading control. Blots were independently replicated with additional extracted total RNA and resulted in similar observations.

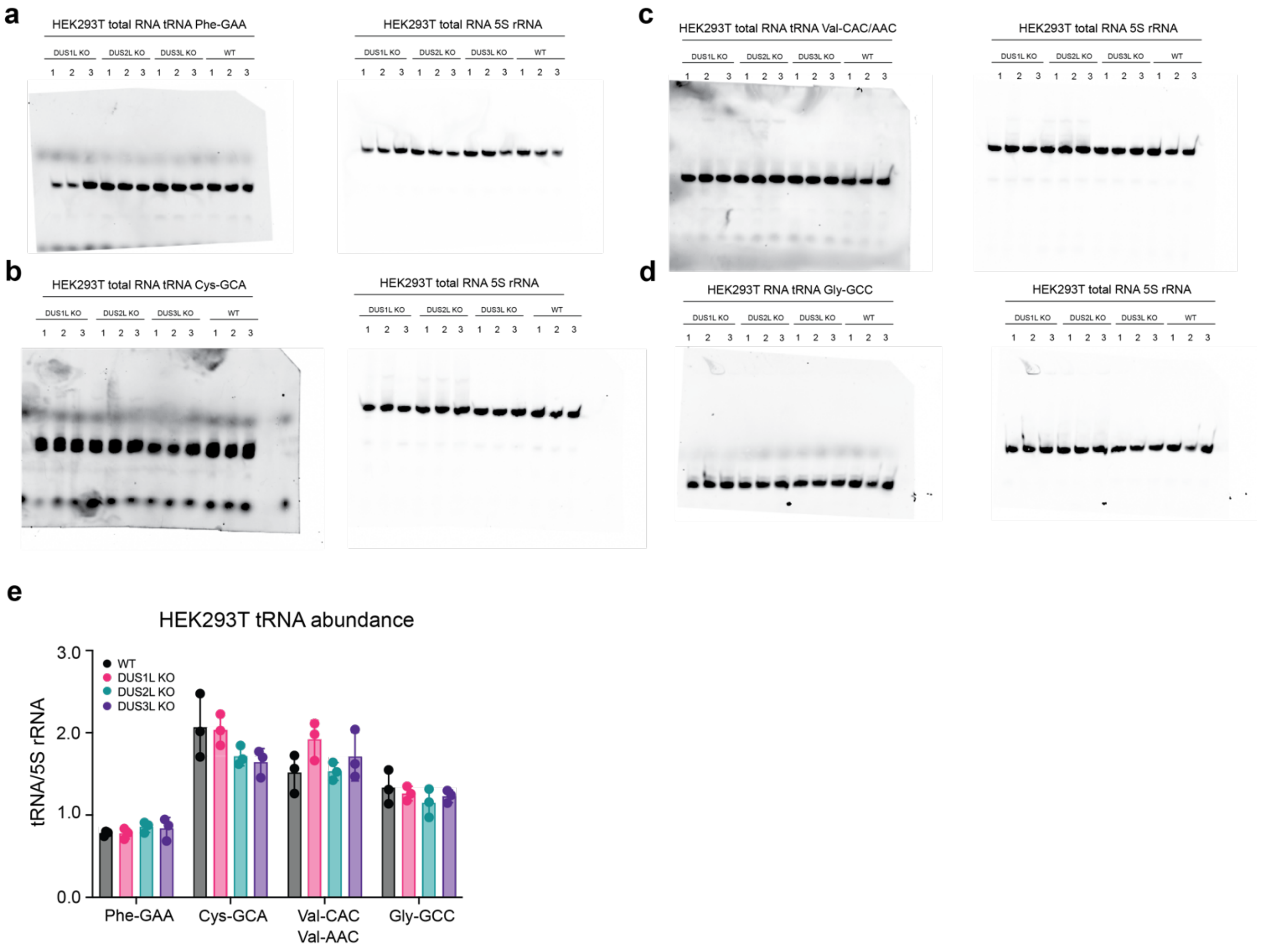


**Supplementary Figure 23.** Northern blot analysis of tRNA abundance in HEK293T cell lines. **(a)** Total RNA from DUS1L KO, DUS2L KO, DUS3L KO, and WT HEK293T cells was probed for tRNA-Phe-GAA using 5S rRNA as a loading control. **(b)** Total RNA from DUS1L KO, DUS2L KO, DUS3L KO, and WT HEK293T cells was probed for tRNA-Cys-GCA using 5S rRNA as a loading control. **(c)** Total RNA from DUS1L KO, DUS2L KO, DUS3L KO, and WT HEK293T cells was probed for tRNA-Val-mAC using 5S rRNA as a loading control. **(d)** Total RNA from DUS1L KO, DUS2L KO, DUS3L KO, and WT HEK293T cells was probed for tRNA-Gly-GCC using 5S rRNA as a loading control. Blots were independently replicated with similar observations. **(e)** Quantification of northern blot analysis of tRNA abundance in HEK293T cell lines. tRNA signal is normalized to 5S rRNA signal from each blot. Three independent biological replicates were analyzed per cell line.

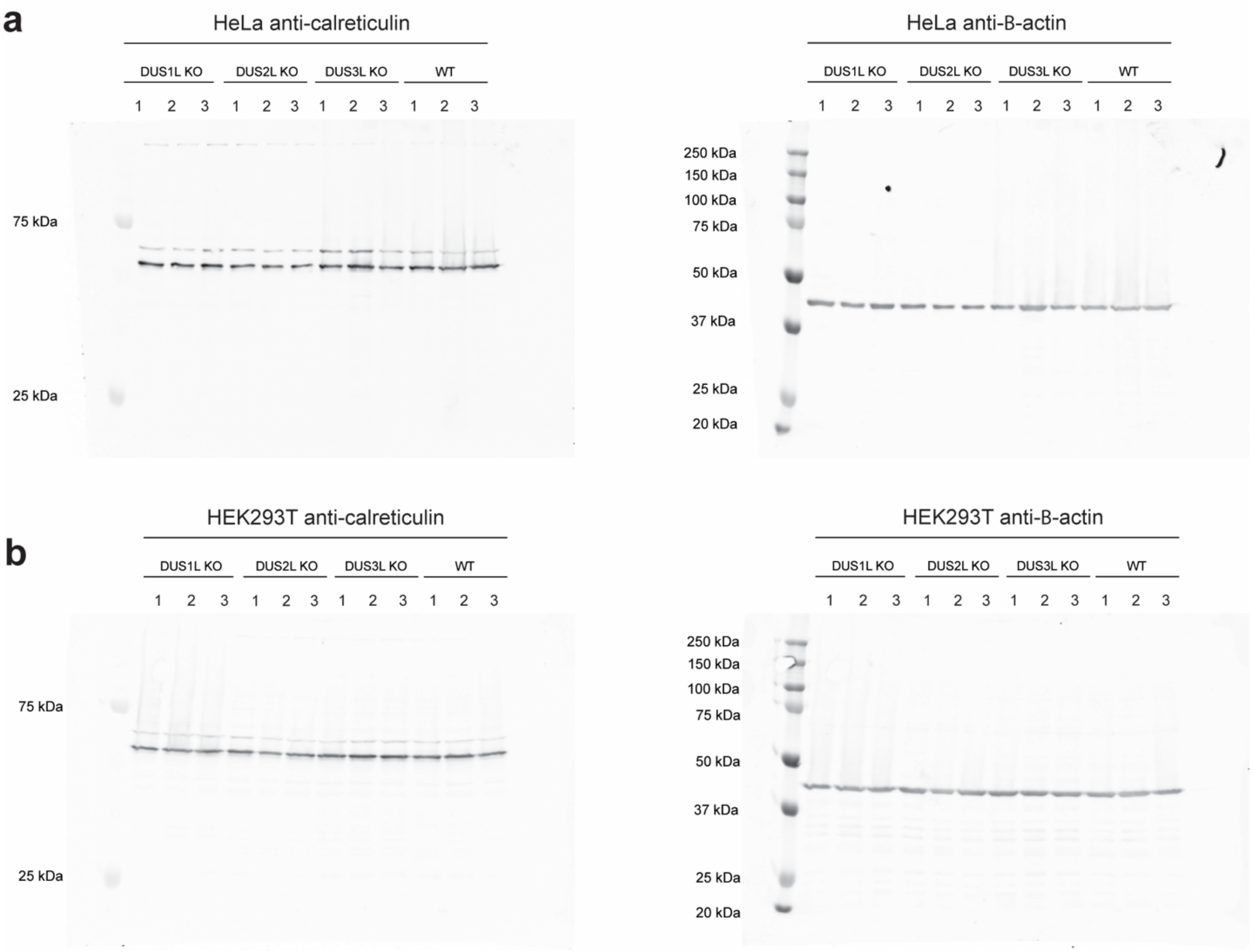


**Supplementary Figure 24.** Western blot analysis of calreticulin expression in HeLa and HEK293T DUS KO cells. **(a)** Western blot of calreticulin in HeLa cell lines; β-actin was used as loading control. **(b)** Western blot of calreticulin in HEK293T cell lines; β-actin was used as loading control. Blots were independently replicated with similar results.
